# Amyloid pathology reduces ELP3 expression and tRNA modifications leading to impaired proteostasis in Alzheimer’s disease models

**DOI:** 10.1101/2023.05.02.538928

**Authors:** Marisa Pereira, Diana R. Ribeiro, Maximilian Berg, Andy P. Tsai, Kwangsik Nho, Stefanie Kaiser, Miguel Moutinho, Ana R. Soares

## Abstract

Alzheimer’s Disease (AD) is a progressive and irreversible neurodegenerative disorder, characterized by the accumulation of abeta-amyloid aggregates, which triggers tau hyperphosphorylation and neuronal loss. While the precise mechanisms underlying neurodegeneration in AD are not entirely understood, it is known that loss of proteostasis is implicated in this process. Maintaining neuronal proteostasis requires proper transfer RNA (tRNA) modifications, which are crucial for optimal translation. However, research into tRNA epitranscriptome in AD is limited, and it is not yet clear how alterations in tRNA modifying enzymes and tRNA modifications might contribute to disease progression. Here, we report that expression of the tRNA modifying enzyme ELP3 is reduced in the brain of AD patients and amyloid AD mouse models, suggesting ELP3 is implicated in proteostasis dysregulation observed in AD. To investigate the role of ELP3 specifically in neuronal proteostasis impairments in the context of amyloid pathology, we analyzed SH-SY5Y neuronal cells carrying the amyloidogenic Swedish familial AD mutation in the APP gene (SH-SWE) or the wild-type gene (SH-WT). Similarly to the amyloid mouse models, SH-SWE exhibited reduced levels of ELP3 which was associated with tRNA hypomodifications and reduced abundance, as well as proteostasis impairments. Furthermore, the knock-down of ELP3 in SH-WT recapitulated the proteostasis impairments observed in SH-SWE cells. Importantly, the correction of tRNA deficits due to ELP3 reduction rescued and reverted proteostasis impairments of SH-SWE and SH-WT knock-down for ELP3, respectively. Additionally, SH-WT exposed to the secretome of SH-SWE or synthetic amyloid aggregates recapitulate the SH-SWE phenotype, characterized by reduced ELP3 expression, tRNA hypomodification and increased protein aggregation. Taken together, our data suggest that amyloid pathology dysregulates neuronal proteostasis through the reduction of ELP3 and tRNA modifications. This study highlights the modulation of tRNA modifications as a potential therapeutic avenue to restore neuronal proteostasis in AD and preserve neuronal function.

**Graphical Abstract:** 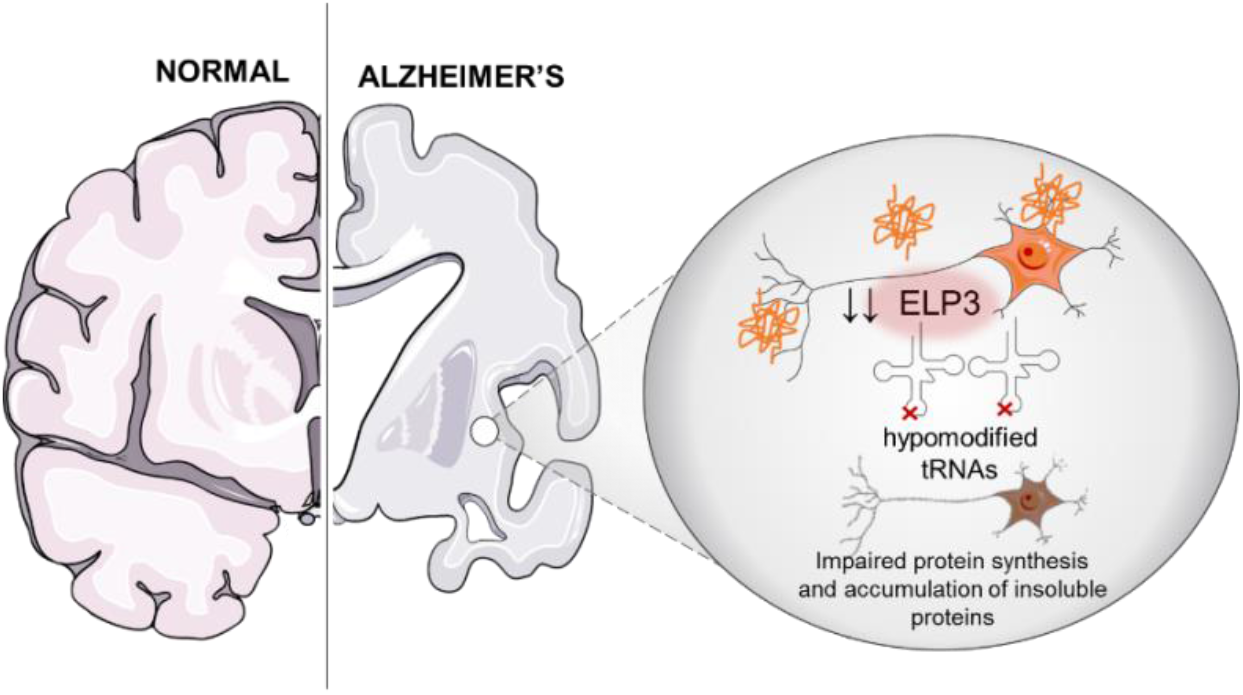

## 1. Introduction

Alzheimer’s disease (AD) is a progressive neurodegenerative disorder and the leading cause of dementia in the elderly (‘Alzheimer’s disease facts and figures’, 2021). It is often characterized by early memory impairment and progressive cognitive decline that results from synapse loss and neuronal atrophy predominately throughout the hippocampus and cerebral cortex (Sheppard and Coleman, 2020). Pathologically, AD is characterized by an abnormal accumulation of protein aggregates, namely extracellular deposition of amyloid-β (Aβ) peptides in senile plaques and the intracellular accumulation of hyperphosphorylated tau (P-tau) that forms neurofibrillary tangles, followed by neurodegeneration (Deture and Dickson, 2019; Sheppard and Coleman, 2020). Despite the underlying mechanisms are still largely unknown, mounting evidence suggests that the accumulation and aggregation of Aβ is a key initiating factor in a cascade of events that lead to this disorder, which include P-tau tangles and neuronal loss (Selkoe and Hardy, 2016). There are also reports indicating that the dissemination of Aβ and P-tau by small extracellular vesicles or exosomes further promotes neurodegeneration (Sardar Sinha *et al*., 2018; Pérez, Avila and Hernández, 2019), and cognitive impairment in AD that has been associated with dysregulation of RNA and protein expression profiles in the brain (Neueder, 2019).

Dysregulation of protein translation is emerging as a central mechanism in the pathogenesis of neurodegenerative disorders (Lee *et al*., 2006; Hanada *et al*., 2013; Ishimura *et al*., 2014; Martin *et al*., 2014; Tuorto and Parlato, 2019). Neurons are particularly dependent on spatial and temporal control of mRNA translation, and disruptions in translation components have a dramatic impact on neuronal survival (Tuorto and Parlato, 2019). One of these components are transfer RNAs (tRNAs) (Phizicky and Hopper, 2010), that recognize mRNA codons through their anticodons to decode the 20 standard amino acids, linking the genetic code information to amino acid identity (Pan, 2018). To be fully active, tRNAs undergo extensive post-transcriptional modifications, catalyzed by different tRNA modifying enzymes (Pereira *et al*., 2018). From all RNA molecules, tRNAs are the most chemically modified class, with an average of 13 modifications per tRNA, that are collectively known as the tRNA epitranscriptome (Pan, 2018). These post-transcriptional chemical modifications are pivotal for tRNA stability and translation efficiency and fidelity (Pan, 2018; Pereira *et al*., 2018). Modifications occurring outside the anticodon loop of tRNAs are crucial for tRNA stability and recognition by aminoacyltransferases. On the other hand, modifications that occur at the anticodon loop, particularly at the wobble position, ensure the diversity of codon recognition and translation efficiency as they optimize mRNA decoding, and improve or stabilize codon-anticodon interactions, contributing to maintain proteome integrity by counteracting protein aggregation (Phizicky and Hopper, 2010; Pan, 2018). Disruptions in tRNA modifications have a direct impact on tRNA expression, mRNA decoding and protein translation, leading to pathological conditions known as “tRNA modopathies” (Chujo and Tomizawa, 2021).

Growing evidence shows that the human brain is particularly sensitive to tRNA epitranscriptome defects (Angelova *et al*., 2018). tRNA modifications, particularly the ones at the anticodon, and their corresponding tRNA modifying enzymes, have been implicated in a panoply of neurological diseases (Pereira *et al*., 2018), namely intellectual disability (Alazami *et al*., 2013; Davarniya *et al*., 2015; Kojic *et al*., 2021), familial dysautonomia (Anderson *et al*., 2001; Lefler *et al*., 2015), and motor neuron diseases such as amyotrophic lateral sclerosis (ALS) (Simpson *et al*., 2009). In general terms, disruption of tRNA modifying enzymes occur in these diseases, affecting the levels of tRNA modifications and leading to defects in mRNA translation in a codon-dependent manner. Quite surprisingly, the impact of tRNA modifications and related networks in AD pathophysiology has been poorly explored and characterized. However, there are indications showing that the tRNA epitranscriptome may represent an AD modifier. For example, a recent report showed m^1^A tRNA hypomodification and decreased expression of the corresponding tRNA modifying enzyme in the AD amyloidogenic 5xFAD mouse model (Shafik *et al*., 2022), and NSUN2, a methyltransferase that methylates cytosine to 5-methylcytosine in tRNAs, was found downregulated in the hippocampus of early onset AD patients (Wu *et al*., 2021). Also, global reduction of protein synthesis rate and increased levels of phosphorylated eIF2α (eIF2α -P) have been found in the cortex and hippocampus of AD patients and mouse models (Vassar, 2009; Ma *et al*., 2013; Stutzbach *et al*., 2014), similarly to what we have observed when inducing tRNA mutations or tRNA hypomodification in vertebrates and in human cell lines (Reverendo *et al*., 2014; Varanda *et al*., 2020; Pereira *et al*., 2021), further re-enforcing the relevance of tRNA molecules and their modifications in the AD context. Compelled by these clues, we hypothesized that dysregulation of tRNA modifying enzymes and, consequently, of tRNA modifications occurs in AD and lead to neuronal proteostasis impairment.

Here, we report that ELP3 expression is reduced in the brain tissue of AD patients and of amyloidogenic 5xFAD mice and in neuronal cell models expressing the *Swedish* familial AD mutation. Our results suggest that amyloid pathology reduces ELP3 expression and consequently leads to tRNA hypomodification, proteostasis impairments and toxicity in neuronal cells. Importantly, we provide evidence that correction of tRNA deficits incurring from ELP3 deficiency rescues proteostasis reinforcing that tRNAs and tRNA modifications are essential to maintain proteostasis in AD and that the tRNA epitranscriptome can represent a promising therapeutic target.

## 2. Results

### 2.1. *ELP3* is differentially expressed in AD patients and its abundance negatively correlates with amyloid plaque density

To evaluate the expression of tRNA modifying enzymes across different human brain regions we used harmonized RNA-seq data generated from three cohort studies, namely, Religious Orders Study and Memory and Aging Project (ROSMAP), Mount Sinai Brain Bank (MSBB), and Mayo RNA-seq (Mayo) available online at *https://agora.adknowledgeportal.org/* (Greenwood *et al*., 2020). We assessed the gene expression of 56 known tRNA modifying enzymes in the different brain regions available on these datasets and found that several were significantly differential expressed (S_Table1). From those, the majority was negatively differentially expressed between AD cases and controls in at least one brain region, particularly the tRNA modifying enzymes that catalyze modifications at the wobble position (Table 1).

**Table 1:**
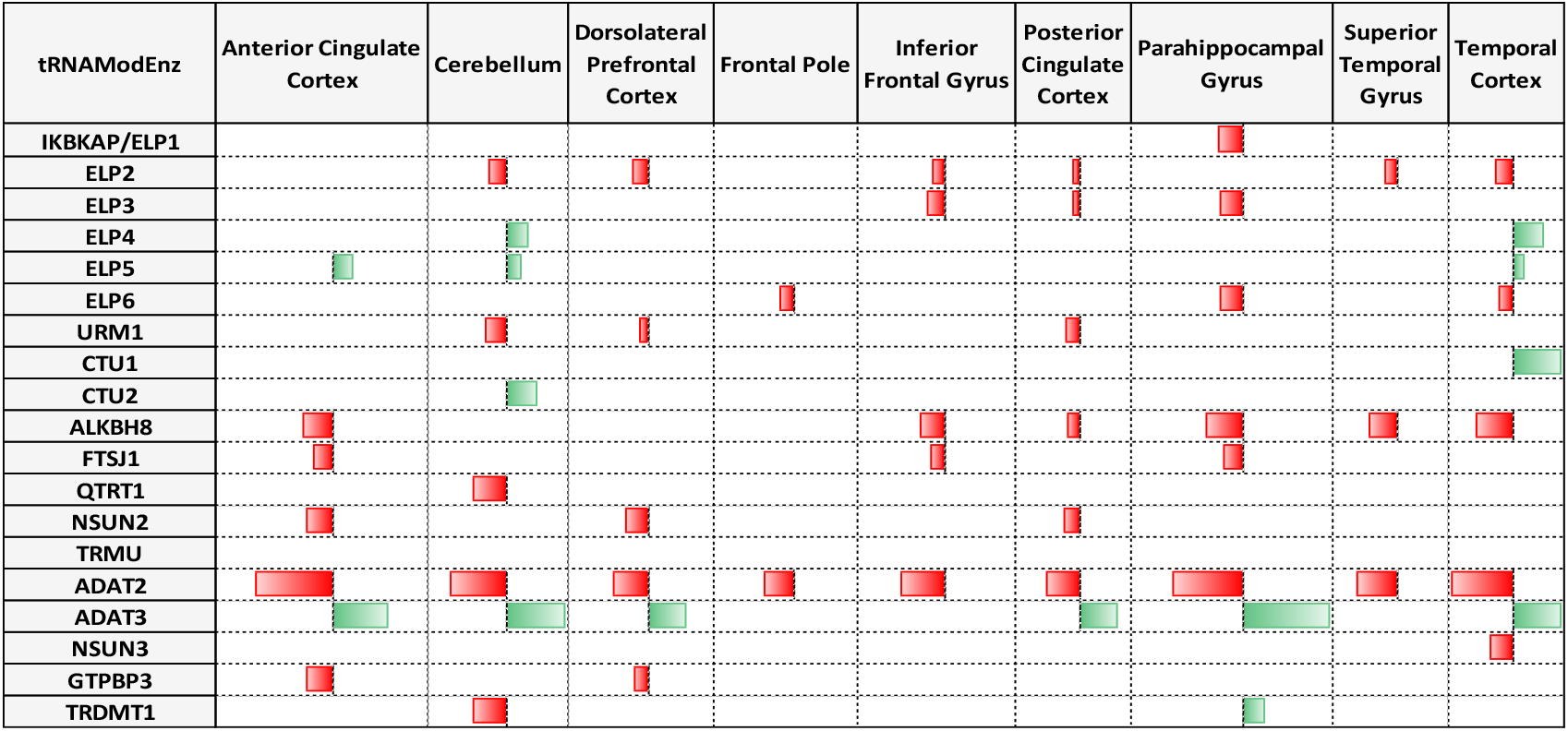
Visual representation of the significant differentially expressed tRNA modifying enzymes that catalyze modifications at the wobble position, in different brain areas of Alzheimer’s disease patients, compared to cognitively normal older adults. Note that red bars indicate negative differential expression levels and green bars indicates positive differential expression values (p-value <0.05). Values range from −4.3E-01 (lowest) to 3.9E-01 (highest). Abbreviations: tRNA modifying enzymes (tRNAModEnz).

Among these we found NSUN2 whose expression was previously found decreased in the hippocampus of early onset AD patients (Wu *et al*., 2021). ADAT2 and ADAT3, that catalyze wobble adenosine-to-inosine modifications, were the 2 enzymes displaying the highest differentially expression, but in a compensatory manner. Whenever ADAT2 was negatively differentially expressed, ADAT3 was positively differentially expressed in a similar magnitude for most of the tissues assessed. Consistently, tRNA modifying enzymes that modify wobble uridine modifications and belong to the elongator complex, namely *ELP1, ELP3* and *ELP6* were negatively differentially expressed in the Parahippocampal Gyrus (PHG), one of the most affected brain regions in AD. We further found a significant negative correlation between *ELP3* expression levels and amyloid plaque burden in the PHG (*β*= −0.006, *p*= 3.75E-03) (Fig 1) (Wang *et al*., 2018). Despite the non-significant results for the Inferior Frontal Gyrus, where *ELP3* is also differentially expressed, the overall tendency observed supports the negative correlation found in the PHG area.

**Figure 1.**
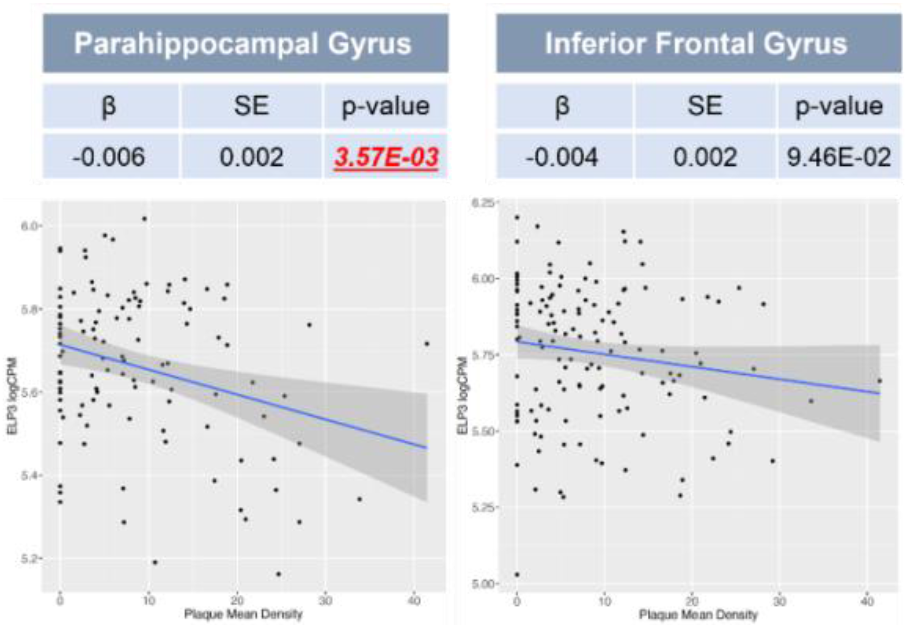
Association between *ELP3* expression and amyloid plaque mean density (number of plaques/mm^2^) in the Parahippocampal Gyrus, and Inferior Frontal Gyrus. The scatter plot shows a significant negative association between ELP3 expression and plaque mean density in the Parahippocampal Gyrus (β= -0.006, *p-*value=3.52E-03). Data extracted from the MSBB cohort (Wang *et al*., 2018). Abbreviations: β coefficient (β) and standard error (SE).

This analysis was complemented with subsequent convergent functional genomic (CFG) analysis that aims to prioritize AD candidate genes, using the AlzData datasets (http://www.alzdata.org/). This revealed that from the differentially expressed tRNA modifying enzymes in AD patients, *ELP3* was the only that cumulative revealed a significant negative correlation between its expression with amyloid and tau pathology based on different amyloidogenic (APPK670N/M671L and PSEN1 M146V mutations) and tauopathy (MAPT P301L mutation) murine models (S_Table2; Table 2), and that was identified as an early differential expressed gene, retrieving the highest CFG score (4 out of 5) for the tRNA modifying enzymes analyzed.

**Table 2:**
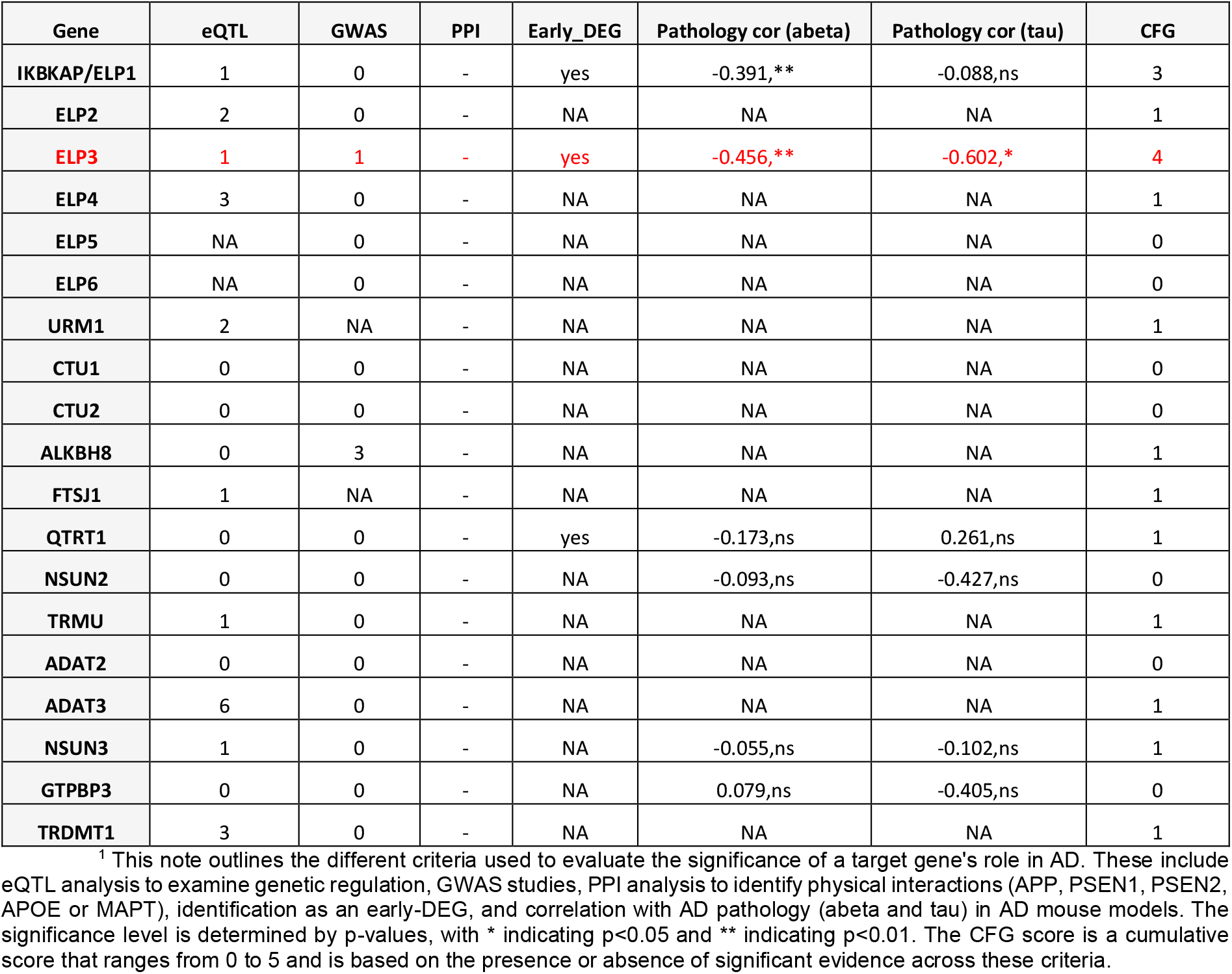
Convergent functional genomic (CFG) analysis for target genes. The *ELP3* gene is highlighted in red. Information collected from http://www.alzdata.org/ ^1^. Abbreviations: expression quantitative trait loci (eQTL); genome-wide association studies (GWAS); protein-protein interaction (PPI), and differentially expressed genes (DEG).

Collectively, this data pointed towards the importance of *ELP3* in amyloid pathology, which led us to focus on this enzyme and its tRNA catalyzed modifications for the subsequent studies.

### 2.2. ELP3 expression and its dependent tRNA modifications are decreased in the hippocampus of 5xFAD mice

Since our patient and animal AD model dataset analysis identified *ELP3* as an early differential expressed gene, we decided to validate this finding in the hippocampus of the amyloid mouse model 5xFAD, that expresses five human familial AD mutations. This model depicts Aβ_1-42_ overproduction, widespread plaque deposition, alterations in immune-related genes, neuronal loss and cognitive decline (Oakley *et al*., 2006). We analyzed *ELP3* mRNA expression in the hippocampus of 1 month old (1M), 4M and 8M 5xFAD mice and C57BL/6J controls (B6) (Fig 2A). We observed a statistically significant decrease of *ELP3* mRNA (∼15% decrease) and protein expression (∼40% decrease) only in the hippocampus of 4M 5xFAD mice (Fig 2B-C), a stage that coincides with learning and memory deficits when amyloid plaques and astrogliosis is increased, which are hallmarks of the prodromal stage of AD, that precedes irreversible degeneration (Girard *et al*., 2014).

**Figure 2:**
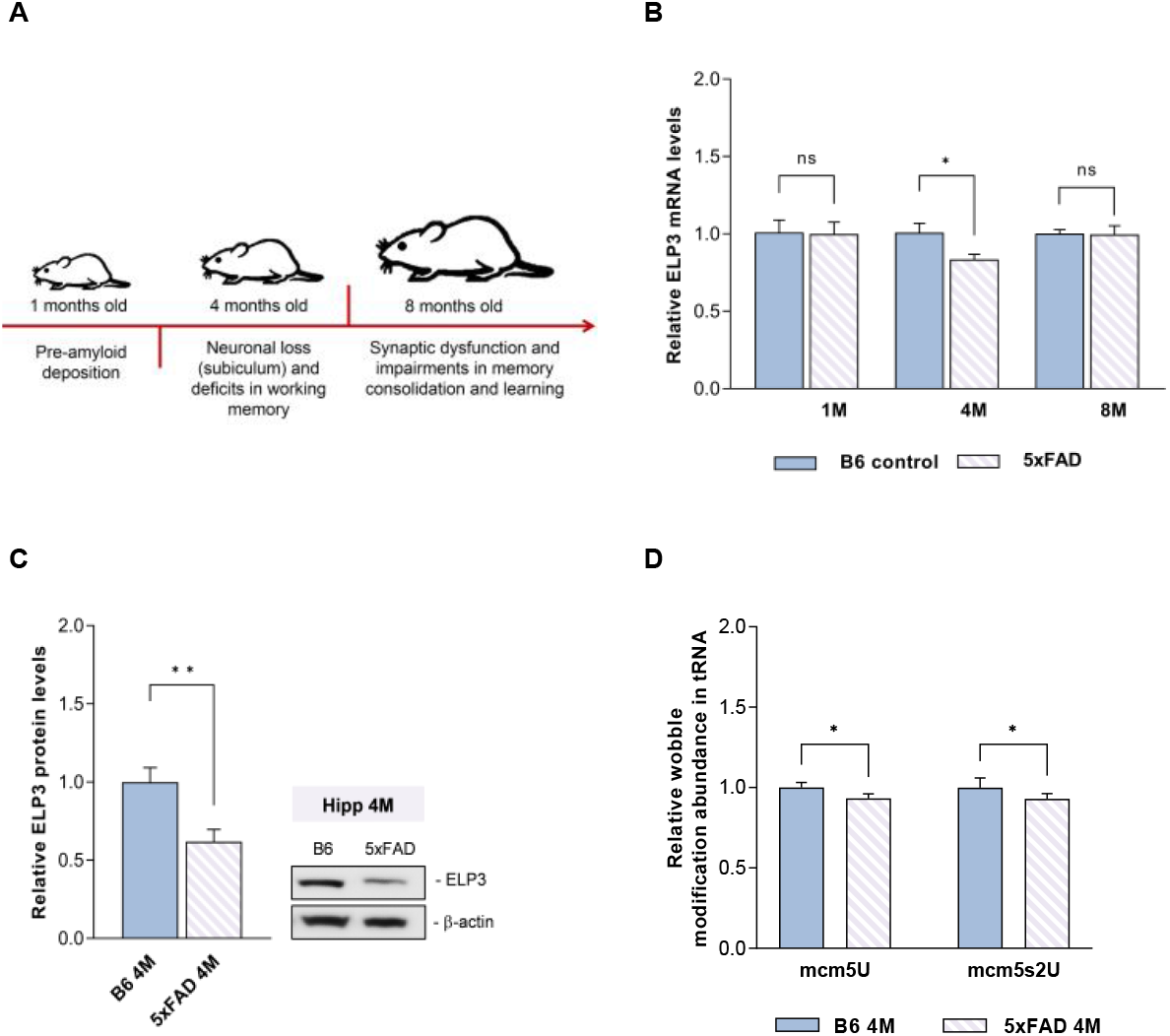
ELP3 is less abundant in the hippocampus of 4 month old (4M) 5xFAD mice. **A)** Schematic representation of the primary pathological alterations in 5xFAD mice at 1, 4, and 8 months old (1M, 4M, and 8M). B) qPCR analysis of ELP3 mRNA levels in the hippocampus (Hipp) of 5xFAD and non-transgenic control (B6) animals. ELP3 mRNA levels were significantly decreased in the Hipp of 5xFAD mice at 4M when compared to the control B6 mice. **C)** Western Blot and graphical representation of ELP3 protein levels in the hippocampus (Hipp) of 4M 5xFAD and non-transgenic control (B6) animals. ELP3 protein levels were significantly decreased in the Hipp of 5xFAD at 4M compared to control B6 mice. β-actin was used as an internal control. **D)** Quantification of the wobble tRNA modifications in the hippocampus (Hipp) of 5xFAD and non-transgenic control (B6) animals at 4M. A significant decrease in the levels of mcm^5^U and mcm^5^s^2^U modifications was detected in the Hipp of 5xFAD at 4M compared to control B6 mice. **Data information**: data are expressed as mean with SEM, n = 3 biological replicates. *p-value <0.05, **p-value <0.01 and non-significant (ns) p-value as assessed by unpaired t test (in B-D).

The Elongator complex and its catalytical subunit ELP3 are essential for cm^5^U catalysis that precedes the mcm^5^U and the final mcm^5^s^2^U modification at the wobble position of three specific tRNAs: tRNA-Glu^UUC^, tRNA-Gln^UUG^ and tRNA-Lys^UUU^ (Abbassi *et al*., 2020). Upon quantification of tRNA modification levels by liquid-chromatography coupled with tandem mass spectrometry (LC-MS/MS) both mcm^5^U and mcm^5^s^2^U modifications were significantly decreased in the hippocampus of 4M 5xFAD mice in comparison to control mice (Fig 2D). Levels of the other tRNA modifications assessed were not affected (Suppl.Fig1-A).

### 2.3. ELP3 abundance is decreased in cellular models bearing the familial AD Swedish mutation

To further understand the implications of *ELP3* differential abundance in the context of AD, we performed different *in vitro* experiments using human neuroblastoma cells (SH-SY5Y) stably expressing either wild-type APP (from now on referred to as SH-WT), or the APP695 mutation, a familial AD mutation, also known as the Swedish mutation (from now on referred to as SH-SWE) (Tao and Wensheng, 2009; Garcia *et al*., 2021). This APP695 mutation occurs at the β-secretase cleavage site, resulting in the production and deposition of toxic amyloid-β (Aβ) forms (Fig 3A) and is also one of the familial AD mutations present in the 5xFAD mice used in this study. Western blotting of APP confirmed its high expression on the SH-WT and SH-SWE cell lines when compared to the original SH-SY5Y cell line, as expected (Fig 3A). Furthermore, according to the Human Protein Atlas (www.proteinatlas.org), ELP3 is expressed at higher levels in neuronal cells than in glial cells in brain tissue, which is consistent with observations in mice (Bento-Abreu *et al*., 2018).

**Figure 3:**
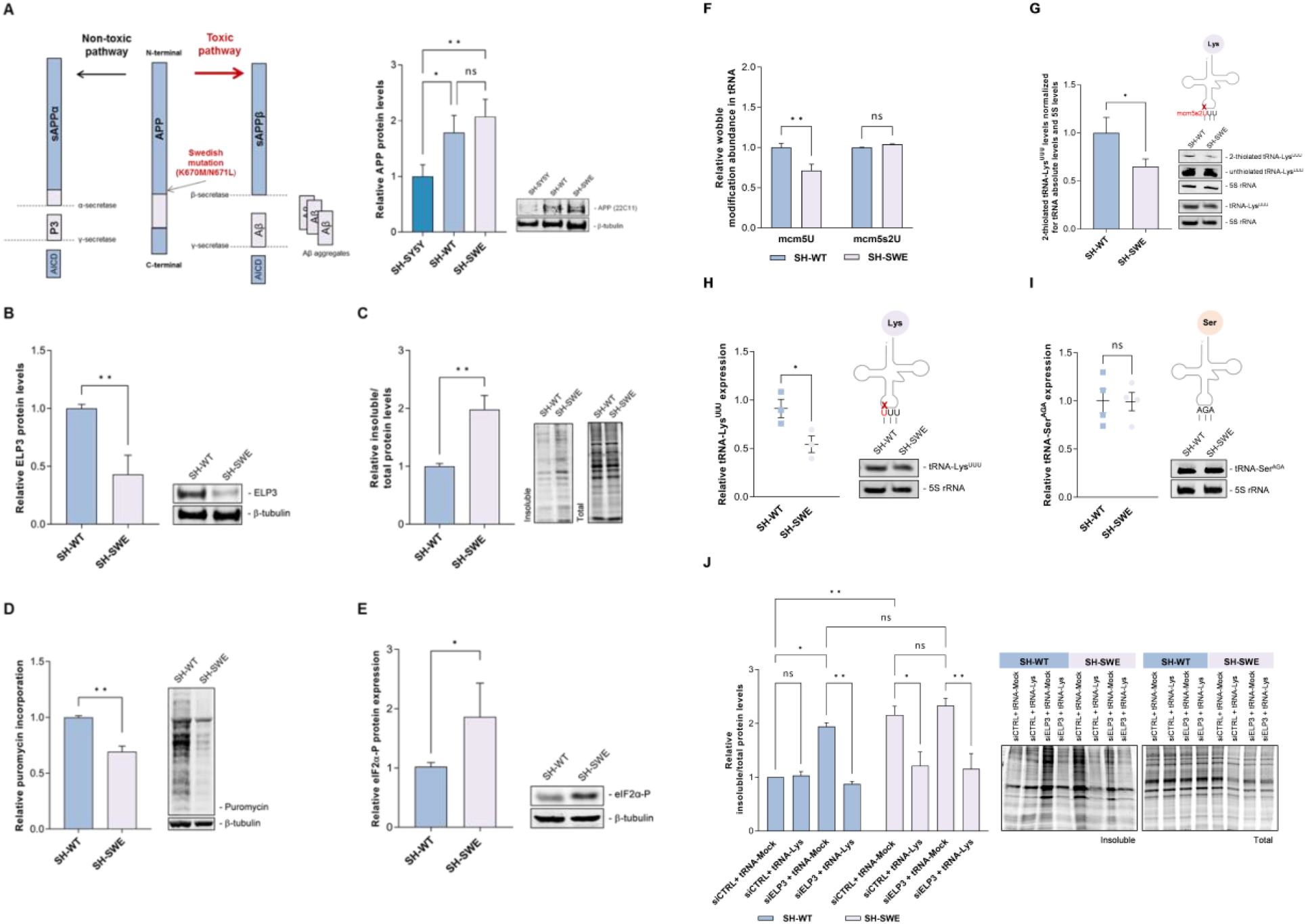
ELP3 expression and ELP3-dependent tRNA modifications are decreased in SH-SWE cells, and protein synthesis is impaired. **A)** Schematic representation of the Swedish mutation in the β-site of APP, which shifts APP processing towards the toxic pathway due to an increase in Aβ production. Western Blot and graphical representation of APP protein levels in SH-SY5Y, SH-WT and SH-SWE cells. A significant increase in APP expression was observed in both SH-WT and SH-SWE cells compared to SH-SY5Y cells. β-tubulin was used as an internal control. **B)** Quantification of ELP3 protein levels by Western blotting. A significant decrease in the expression of ELP3 was observed in SH-SWE cells when compared to SH-WT cells. βtubulin was used as an internal control. **C)** Quantification of relative insoluble protein fraction. A significant increase in the amount of insoluble protein levels was observed in SH-SWE cells compared to SH-WT cells. Representative acrylamide gels of both insoluble and total fractions after blue safe staining are depicted. **D)** Relative rate of protein synthesis based on the incorporation of puromycin by the SUNSet method. A significant decrease in puromycin incorporation during translation was observed in SH-SWE cells compared to SH-WT cells by Western blotting. β-tubulin was used as an internal control. **E)** Quantification of eIF2α-P expression by Western blotting. A significant increase in eIF2α-P was observed in SH-SWE cells compared to SH-WT cells. β-tubulin was used as an internal control. **F)** Quantification of the U34 tRNA modifications in SH-SWE cells by LC/MS-MS. A significant decrease in the level of the ELP3 dependent mcm^5^U modification was observed in SH-SWE cells compared to SH-WT cells. **G)** Quantification of 2-thiolated tRNA-Lys^UUU^ levels by Northern blotting. A significant decrease in the abundance of 2-thiolated tRNA-Lys^UUU^ in SH-SWE cells was observed in comparison with SH-WT cells, after normalizing for the total tRNA-Lys^UUU^. 5S rRNA was used as internal control. **H)** Quantification of mature tRNA-Lys^UUU^ abundance by Northern blotting. Abundance of mature tRNA-Lys^UUU^ was significantly decreased SH-SWE cells, when compared to SH-WT cells. 5S rRNA was used as internal control. **I)** Quantification of tRNA-Ser^AGA^ abundance by Northern blotting. As expected, no significant alterations in the abundance of the total tRNA-Ser^AGA^ was observed between SH-SWE and SH-WT cells. 5S rRNA was used as internal control. **J)** Relative insoluble protein fraction quantifications after transfection of a tRNA-Lys^UUU^ plasmid in ELP3 silenced SH-WT and SH-SWE cells, followed by a representative acrylamide gel staining with blue safe. Cells were co-transfected with the siCTRL and tRNA-Mock plasmid (siCTRL+ tRNA-Mock), siCTRL and tRNA-Lys^UUU^ plasmid (siCTRL+ tRNA-Lys), siRNA against ELP3 and tRNA-Mock plasmid (siELP3 + tRNA-Mock), or siRNA against ELP3 and tRNA-Lys^UUU^ plasmid (siELP3 + tRNA-Lys). Silencing of ELP3 in SH-WT cells resulted in a significant increase in the abundance of insoluble proteins, which was restored upon co-transfection with the tRNA-Lys^UUU^ plasmid. The insoluble protein fraction was also significantly increased in both siCTRL and siELP3-transfected SH-SWE cells, and was restored, in both situations, upon co-transfection with the tRNA-Lys plasmid.Data information: data are expressed as mean with SEM, n = 3 biological replicates. *p-value <0.05, **p-value <0.01, and non-significant (ns) p-value as assessed by two-way ANOVA with the Sidak test (in A, E and I), and unpaired t test (in B-D, and F-H).

ELP3 protein expression was significantly decreased in SH-SWE cells compared to SH-WT cells, by approximately 50% (Fig 3B). Additionally, the expression of ELP3 was maintained between the undifferentiated and differentiated state in both cell lines (Suppl.Fig2-A), suggesting that the levels of ELP3 do not change with differentiation, but change in the presence of the familial SWE mutation. SH-SWE cells were also characterized by significant accumulation of insoluble proteins (Fig 3C) and impaired protein synthesis, as detected by the puromycin incorporation assay SUnSET (Schmidt *et al*., 2009) (Fig 3D), and increased eIF2α-P (Fig 3E). This was expected since APP695 mutation leads to deposition of toxic Aβ forms and decreased ELP3 levels are associated with translation impairments through increased levels of eIF2α-P and accumulation of protein aggregates as demonstrated by us and others (Endres, Dedon and Begley, 2015; Pollo-Oliveira *et al*., 2020; Tavares *et al*., 2021).

The lower expression of ELP3 in SH-SWE cells was also accompanied by a significant decrease in mcm^5^U levels (Fig 3F), but not in mcm^5^s^2^U levels (Fig 3F). As this was unexpected, we analyze the thiolation levels of tRNA-Lys^UUU^, one of the 3 tRNAs that carries the final mcm^5^s^2^U modification, by an alternative method that consists on an APM gel electrophoresis, combined with Northern blotting for the tRNA of interest (Igloi, 1988; Lemau de Talancé *et al*., 2011). 2-thiolated tRNAs migrate more slowly compared to non-thiolated tRNAs in the APM gel, which allows to conclude on the amount of tRNA that is 2-thiolated. Although quantification of mcm^5^s^2^U levels in total tRNA by LC/MS-MS did not reflect any alteration between cell lines, we observed a significant decrease in the amount of 2-thiolated tRNAs-Lys^UUU^ in the SH-SWE cells when compared with SH-WT cells (Fig 3G). Even though tRNA-Lys^UUU^ was significantly decreased in SH-SWE cells (Fig 3H), decreased thiolation was not due to a decrease in the tRNA absolute expression as data was normalized for the internal control 5S and for the tRNA-Lys^UUU^ quantity detected by regular Northern blotting (Fig 3H). No differences were detected in any of the other modifications analyzed (Suppl.Fig2-B).

As it is well established that ELP3 and the Elongator complex participates in the final mcm^5^s^2^U modification of 2 other tRNAs, namely tRNA-Glu^UUC^ and tRNA-Gln^UUG^ and that tRNA hypomodification may impact tRNA abundance (Rezgui *et al*., 2013), we have also quantified the expression of the mentioned tRNAs by Northern Blotting. Our data shows that, similarly to tRNA-Lys^UUU^, tRNA-Glu^UUC^ is also significantly decreased in SH-SWE cells (Fig Suppl2-C), but no difference was detected in tRNA-Gln^UUG^ (Fig Suppl2-D), nor in tRNA-Ser^AGA^, which was used as a control (Fig 3I).

To further confirm that decreased ELP3 expression levels in SH-SWE were indeed affecting tRNA modification levels, we took advantage of a plasmid encoding unmodified tRNA-Lys^UUU^. Transfection of this plasmid in SH-SWE cells restored proteostasis by decreasing the accumulation of protein aggregates and reverted its levels to SH-WT insoluble protein fraction levels (Fig 3J). No alterations were found when we overexpressed the tRNA-Lys^UUU^ in SH-WT cells (Fig 3J). Moreover, silencing of ELP3 expression in SH-WT cells (Suppl.Fig3-A,B), mimics the accumulation of insoluble proteins (Fig 3J), and negatively affected tRNA-Lys^UUU^ expression (Suppl.Fig3-C) as observed in SH-SWE cells (Fig 3H). No alterations were detected in the expression of tRNA-Ser^AGA^ (Suppl.Fig3-D). Overexpression of unmodified tRNA-Lys^UUU^ in ELP3 silenced SH-WT also reverted the accumulation of insoluble proteins to basal levels (Fig 3J).

Of note, silencing of ELP3 in SH-SWE cells did not significantly alter the insoluble proteins levels compared to SH-SWE siCTRL probably due to the already reduced levels of ELP3 expression in the SH-SWE cell line. Transfection of tRNA-Lys^UUU^ plasmid in both ELP3 silenced cell lines restored proteostasis by decreasing the accumulation of protein aggregates to basal levels (Fig 3J).

### 2.4. The secretome of SH-SWE cells negatively affects ELP3 abundance and tRNA modification levels and increases the accumulation of insoluble proteins in SH-WT cells

Since SH-SWE cells are characterized by a specific APP mutation that induces accumulation of toxic Aβ forms, we wondered if the observed disruption in ELP3 abundance occurred in response to the proteotoxic stress that is already triggered by the mutation. To elucidate if this was the case, we evaluated the effects produced by the SH-SWE secretome, that includes the secreted toxic Aβ forms, on SH-WT cells. Upon 72h incubation of SH-WT with SH-SWE secretome, ELP3 expression was decreased by ∼30% when compared with SH-WT cells incubated with their own secretome (Fig 4A). These cells recapitulated all SH-SWE findings reported above (Fig 3). There was a significant increase (90% increase) in accumulation of cellular insoluble proteins (Fig 4B), and a decrease in protein synthesis of approximately 15% (Fig 4C). Additionally, mcm^5^U modification levels were negatively impacted in SH-WT cells incubated with SH-SWE secretome as quantified by LC-MS/MS (Fig 4D, Suppl.Fig4-A) and the 2-thiolated tRNA-Lys^UUU^ levels were also reduced as quantified by APM gel electrophoresis combined with Northern blotting (Fig 4E). The tRNA-Lys^UUU^ levels were also decreased after incubation of SH-WT cells with SH-SWE cells secretome (Fig 4F), and no alterations were detected in the expression of tRNA-Glu^UUC^ and tRNA-Gln^UUG^ (Suppl.Fig4-B,C), and tRNA-Ser^AGA^ (Fig 4G). To understand if indeed toxic Aβ fibrils trigger ELP3 dysregulation, we incubated SH-WT cells with pre-aggregated Aβ _1-42_ peptides. This resulted in a ∼50% decrease in ELP3 abundance in SH-WT cells (Fig 4H), confirming that exposure to toxic Aβ aggregates induces a cellular response that triggers tRNA epitranscriptome reprograming through ELP3 expression modulation.

**Figure 4:**
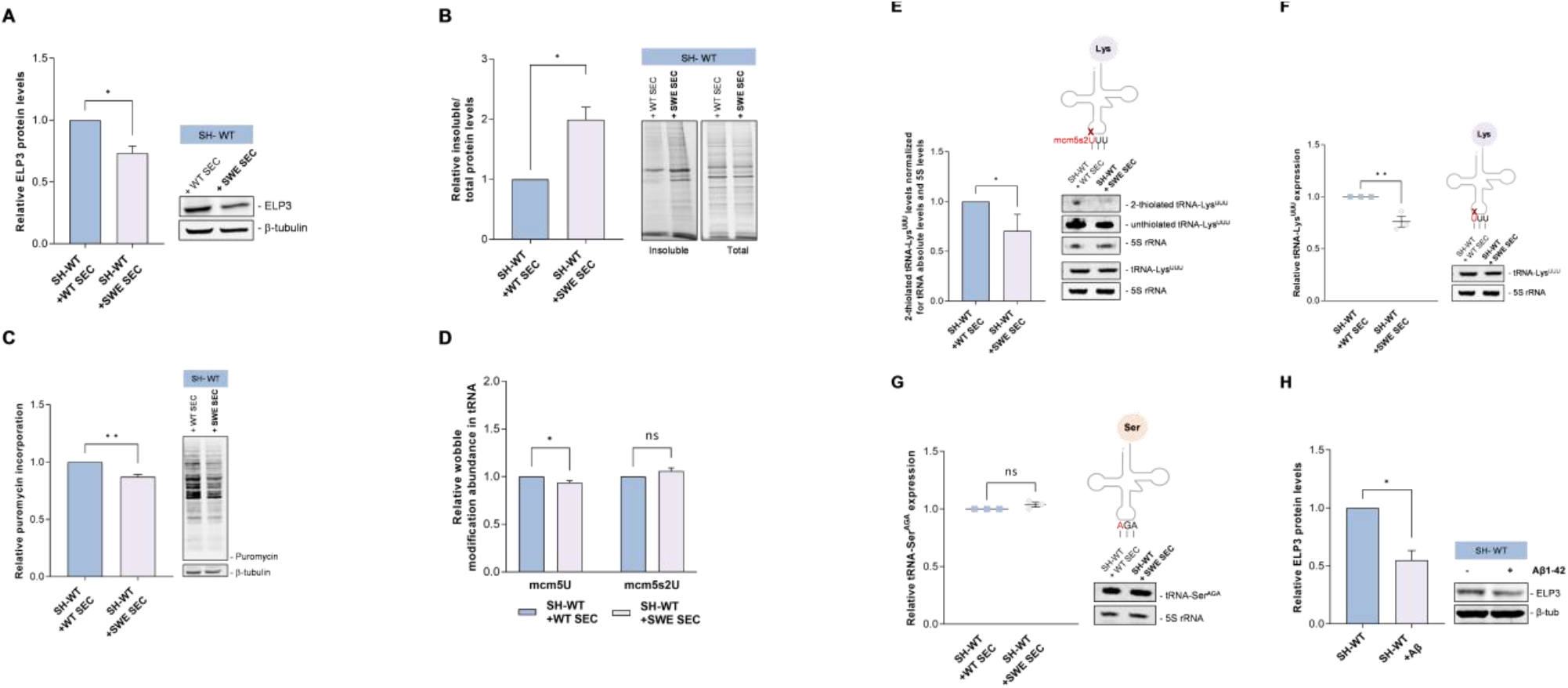
Exposure to the secretome of SH-SWE neuronal cells reduces ELP3 expression and increases the accumulation of insoluble proteins in SH-WT cells. **A)** Quantification of ELP3 protein expression by Western blotting. A significant decrease in the expression of ELP3 was observed in SH-WT cells incubated with SH-SWE secretome (SH-WT + SWE SEC) when compared to SH-WT cells incubated with their own secretome (SH-WT + WT SEC). β-tubulin was used as an internal control. **B)** Quantification of the insoluble protein fraction in SH-WT cells after incubation with SH-SWE secretome (SH-WT + SWE SEC), or with SH-WT secretome (SH-WT + WT SEC). A significant increase in the insoluble protein fraction was observed in SH-WT + SWE SEC cells when compared to SH-WT + WT SEC cells. Representative acrylamide gel after blue safe staining is depicted. **C)** Quantification of protein synthesis rate in SH-WT cells upon incubation with SH-SWE secretome (SH-WT + SWE SEC), or incubation with SH-WT secretome (SH-WT + WT SEC). A significant decrease in protein synthesis rate, identified by a decrease in the incorporation of puromycin, was observed in SH-WT + SWE SEC cells when compared to SH-WT + WT SEC cells. β-tubulin was used as an internal control. **D)** Quantification of ELP3-dependent U34 tRNA modifications. A significant decrease in the levels mcm5U modification was observed in SH-WT + SWE SEC cells when compared to SH-WT + WT SEC cells, in accordance with the decreased levels of ELP3 expression observed. **E)** Quantification of 2-thiolated tRNA-Lys^UUU^ abundance. A significant decrease in the abundance of 2-thiolated tRNA-Lys^UUU^ levels occurs in SH-WT + SWE SEC cells, when compared to SH-WT + WT SEC cells, and normalized for the total tRNA-Lys^UUU^ levels. 5S rRNA was used as internal control. **F)** Quantification of mature tRNA-Lys^UUU^ levels by Northern blotting. A significant decrease in tRNA-Lys^UUU^ abundance is observed after incubation of SH-WT cells with SH-SWE secretome (SH-WT + SWE SEC). 5S rRNA was used as internal control. **G)** Quantification of tRNA-Ser^AGA^ abundance, used as a control tRNA, by Northern blotting. As expected, no significant alterations in the abundance of mature tRNA-Ser^AGA^ was detected between SH-WT + SWE SEC cells, and SH-WT + WT SEC cells. 5S rRNA was used as internal control. **H)** Quantification of ELP3 protein expression levels in SH-WT cells incubated with Aβ1-42 peptides (SH-WT + Aβ). A significant decrease in the expression of ELP3 was observed in SH-WT + Aβ cells when compared to SH-WT cells not exposed to synthetic Aβ. β-tubulin was used as an internal control. **Data information**: data are expressed as mean with SEM, n = 3 biological replicates. *p-value <0.05, **p-value <0.01, and non-significant (ns) p-value as assessed by unpaired t-test (in A-C, E-H), and two-way ANOVA with the Sidak test (in D).

## 3. Discussion

It is now evident that tRNA epitranscriptome dysregulation occurs in a panoply of neurological disorders, as reviewed in (Blanco *et al*., 2014; Pereira *et al*., 2018; Ramos and Fu, 2019). However the reports concerning the relevance of the tRNA epitranscriptome in AD are scarce (Shafik *et al*., 2022), and the role of wobble uridine modifications in neuronal proteotoxic stress generation and their contribution in AD pathogenesis and progression was not yet explored.

Here, we found evidence of dysregulation of the tRNA modifying enzyme *ELP3* in the hippocampus and cortex of AD patients and establish a negative correlation between *ELP3* expression and amyloid severity in the Parahippocampal region (PHG). Previous work has shown that PHG undergoes substantial transcriptomics alterations in AD patients (Neff *et al*., 2021) and is associated with early-Aβ deposition and memory loss events (Wang *et al*., 2016), supporting a link between the severity of the amyloid pathology and ELP3 dysregulation. In line with these findings, we also uncovered the potential role of *ELP3* as an early differential expression gene in different amyloidogenic murine models.

We further confirmed that ELP3 mRNA and protein expression is less abundant in the hippocampus of 4M 5xFAD mice. Different studies reported the existence of extracellular neuronal amyloid deposits and gliosis at this time point that culminate in cognitive deficits, characteristic of the earliest symptomatic stage (Landel *et al*., 2014; Forner *et al*., 2021). As expected, and coinciding with ELP3 decreased expression, at this stage we also detected a decrease in the levels of ELP3-dependent tRNA modifications mcm^5^U and mcm^5^s^2^U.

Similar findings were observed in the *in vitro* AD model SH-SWE cell line that is characterized by mutated APP and increased Aβ production and accumulation, in comparison with SH-WT cells (Fernandes *et al*., 2018). Moreover, we observed that SH-SWE cells are characterized by a generalized increase in accumulation of insoluble proteins and decreased protein synthesis. Remarkably, the accumulation of insoluble proteins in SH-SWE cells was reverted by overexpression of unmodified tRNA-Lys^UUU^, further reinforcing that ELP3 decreased expression in SH-SWE had a direct impact on tRNA modification levels and proteostasis and that lack of ELP3 triggers the accumulation of insoluble proteins due to tRNA-Lys^UUU^ hypomodification. These findings are in line with what was previously shown in yeast lacking ELP3 (Endres, Dedon and Begley, 2015; Pollo-Oliveira *et al*., 2020; Tavares *et al*., 2021), where proteostasis impairments and accumulation of insoluble proteins occur due to tRNA wobble uridine hypomodification. Indeed, the mcm^5^s^2^U34 modification at tRNA-Lys^UUU^ increases the network of interactions to stabilize codon:anticodon base-pairing, contributing to translation efficiency and fidelity (Björk *et al*., 2007; Klassen *et al*., 2015). In addition, studies in yeast strains lacking mcm^5^s^2^U modifications reveal the occurrence of codon-specific translational pausing (Nedialkova and Leidel, 2015; Tavares *et al*., 2021), that increases the accumulation of misfolded proteins and amino acid misincorporations (Tavares *et al*., 2021).

Since we also have an increase in the phosphorylation of eIF2α and a slightly decrease in the protein synthesis rate, we suggest the activation of the integrated stress response (ISR) response which can assure minimal protein synthesis as a protective response. These results are consistent with the increase in eIF2α phosphorylation found in brains of AD patients and in AD mice models (Kim *et al*., 2007; Oliveira *et al*., 2021). Surprisingly, some studies reported that high levels of phosphorylated eIF2α can also increase the expression of β-secretase (BACE1), which accelerate the beta-amyloidogenic pathway involved in the AD pathology (O’Connor *et al*., 2008; Ma *et al*., 2013).

Silencing ELP3 in SH-WT cells caused a similar proteotoxic phenotype as observed in SH-SWE cells, which was rescued by transfection of unmodified tRNA-Lys^UUU^. This points towards the potential deleterious effect of ELP3 dysregulation in neuronal proteostasis. However, ELP3 decreased expression detected in the different AD models tested, does not explain on its own the proteostasis impairments observed in the disease. This is particularly true in the case of the SH-SWE cells that carry a familial AD mutation that in itself contributes to proteotoxic stress due to accumulation of toxic Aβ forms, as previous studies have reported an increase in Aβ40 and Aβ42 secretion in cell culture media from SH-SWE cells compared to SH-WT (Belyaev *et al*., 2010; Jämsä *et al*., 2011). Our results led us to question whether ELP3 disruption in AD patients and AD models carrying the APP695 mutation is a cellular attempt to counteract the proteotoxic stress by decreasing translation rate. After incubation of SH-WT cells with SH-SWE secretome, there was a clear and significant decrease in ELP3 abundance. These cells also depicted a decrease in mcm^5^U34 levels, an increase in the insoluble protein fractions and a decrease in protein synthesis rate (Fig 4), fully recapitulating what occurs in SH-SWE cells. Furthermore, incubation of SH-WT cells with pre-aggregated human amyloid 1-42 also led to a decrease in ELP3 expression, recapitulating the ELP3 levels found in SH-SWE cells. These results suggest that cells alter their tRNA epitranscriptome in the presence of amyloid aggregates, possibly to counteract proteotoxic stress, eliciting the ISR that leads to translation attenuation to allow cells to cope with stressful agents. In fact, a reduction in translation levels has been shown to improve the capacity of the proteostasis network to eliminate aberrant proteins and promote health and longevity (Jiménez-Saucedo, Berlanga and Rodríguez-Gabriel, 2021; Martinez-Miguel *et al*., 2021). However, in this pathological state, the decrease in ELP3 and the consequent tRNA hypomodification seems to further contribute to exacerbate the AD phenotype and is not beneficial for the cell. Indeed, AD is characterized by activation of the ISR and reversal of ISR-mediated translation reprogramming results in neuroprotection (Costa-Mattioli and Walter, 2020), which demonstrates that ISR activation is not beneficial in AD. The fact that proteostasis is recovered by tRNA overexpression also indicates that tRNA hypomodification induced by decreased ELP3 expression is not beneficial in the AD context and further highlights the potential of tRNAs to be used as therapeutics to restore proteostasis in diseases characterized by proteotoxic stress, ISR activation or epitranscriptome disruptions.

Altogether, our results indicate that ELP3 expression is dependent on the levels of aberrant proteins and that modulation of ELP3 expression leads to reprograming of tRNA modifications, with a direct impact on proteostasis and protein synthesis rate. In conclusion, our findings show that the tRNA epitranscriptome is modulated in AD and that this modulation has a direct impact on proteostasis imbalances observed. Thus, developing strategies to reprogram tRNA epitranscriptome in AD will likely increase the probability of recovering corrupted translation and improve neuronal survival, which should be tested soon in different AD models.

## 4. Material and Methods

### 4.1. RNA-Seq analysis of post-mortem human brains

To evaluate the differential expression of ELP3 gene between AD cases and controls we accessed the harmonized RNA-seq data from the National Institute on Aging’s Accelerating Medicines Partnership in AD (AMP-AD) Consortium and the AD research community and available online at Agora Platform (**https://agora.ampadportal.org**).

To investigate the association between ELP3 expression levels and amyloid plaque density (number of plaques/mm^2^) in the PHG brain region, we used linear regression models (R version 3.6.1).

### 4.2. 5xFAD Mouse Model

5xFAD mice were obtained from the Jackson Laboratory (stock no. 34840-JAX) and express five human familial AD mutations driven by the mouse Thy1 promoter (APPSwFlLon, PSEN1*M146L* L286V]6799Vas). To collect brain tissue, mice were anesthetized with 1.2% 2,2,2-tribromoethanol (Avertin), perfused with ice-cold phosphate buffered saline (PBS), and the brain removed, dissected, and stored at -80°C until further use. Animal experiments were conducted under the approved Indiana University School of Medicine Institutional Animal Care and Use Committee (IACUC) protocol number 22115.

For qPCR and Western blot analysis, tissue was homogenized in buffer containing 1% NP-40, 0.5% sodium deoxycholate, 0.1% SDS and protease inhibitor cocktail (Sigma Aldrich, #P8340). For RNA extraction, tissue homogenate was mixed in an equal volume of RNA-Bee (Amsbio, CS-104B) and RNA was isolated using phenol-chloroform extraction and a Purelink RNA Mini Kit (Life Technologies, 12183020) with an on-column DNAse Purelink Kit (Life Technologies, 12183025). 1000-500 ng of RNA were converted to cDNA with the High-Capacity RNA-to-cDNA kit (Applied biosystems, 4388950) and qPCR was performed on StepOne Real Time PCR System with Taqman Assays (Life Technologies). The mRNA expression of *Elp3* (Taqman assay Mm00804536_m1) was normalized to *Gapdh* (Taqman assay Mm99999915_g1), and expressed as fold changes relative to controls, using the ΔΔCt method. Protein extracts were obtained by centrifugation of the tissue homogenate 12,000 g at 4 °C for 15 minutes, and recovery of the supernatant. For Western blot, protein extracts were heated for 5 min at 95 °C, loaded into 4–12% Bis-Tris gels (Life Technologies), and run at 150V. Proteins were transferred into immobilon-PPVDF membranes at 400 mA, blocked in 5% milk in TBS-Tween 0.1%, and incubated with primary antibodies overnight at 4 °C. All secondary HRP-conjugated antibodies were incubated for 1 h at room temperature. The following primary antibodies were used: Elp3 (Cell Signaling 5728S) and β-actin (Santa Cruz sc-517582).

### 4.3. Cell culture and siRNA transfections

SH-SY5Y-WT and SH-SY5Y-appSwe were cultured in Dulbecco’s Modified Eagle Medium (DMEM, Gibco, Cat.11965084), supplemented with 10% of Fetal Bovine Serum (FBS, Sigma-Aldrich, Cat.F1051,) and 2% of Pen-Strep-Glut (Gibco, Cat.15070063).

siGenome SMARTpool human siRNA targeting ELP3 (siELP3), and a negative control siRNA (siCTRL) were transfected into the SH-SY5Y cells in 24 well plates. Briefly, 10 nM of each siRNA were diluted in Opti-MEM (Gibco, Cat.31985062), followed by addition of 1 μL/well of DharmaFECT 1 (Dharmacon, Cat.T-2001-02). After an incubation of 20 minutes, 4×10^4^ cells/mL were added to each well in DMEM supplemented with 10% FBS and allowed to growth 72 hours at 37 °C in a CO^2^ incubator.

### 4.4. Total RNA extraction

The total RNA from cell culture and mouse tissues were extracted using the TRIsure™ (Bioline, Cat.BIO-38033) reagent according to manufacturer’s instructions.

In the case of RNA extraction from tissues, 50-100 mg were first homogenized in a glass homogenizer (TissueRuptor II - Qiagen, Cat.9001272).

The integrity and quantity of each RNA sample were assessed using a spectrophotometer DeNovix DS-11 and the the 4200 TapeStation System or a Bioanalyzer.

### 4.5. LC-MS/MS of tRNA modification abundance

Transfer RNAs (tRNAs) were isolated from total RNA by size exclusion chromatography (SEC), using an Agilent SEC 300 Å column (Agilent 1100 HPLC system) and ammoniumacetate (0.1 M, pH = 7) as mobile phase at 1mL/min and 40°C. The collected tRNAs were vacuum concentrated (Speedvac, Thermo Fisher Scientific) and precipitated at -20 °C overnight after adding 0.1× vol. ammoniumacetate (5 M) and 2.5× vol. ethanol (100%). After centrifugation at 12.000xg for 30 minutes at 4 °C, the resulting tRNA pellets were washed with 70% ethanol and resuspended in pure water. Following UV quantification at 260 nm (NP80, IMPLEN), 200 ng of the purified tRNAs were digested into nucleosides using benzonase (2 U), phosphodiesterase I (0.2 U), alkaline phosphatase (2 U), Tris pH 8.0 (5 mM), magnesium chloride (1 mM), tetrahydrouridine (5 μg), butylated hydroxytoluene (10 μM) and pentostatin (1 μg) in a final volume of 20 μL. Upon a 2 h incubation at 37 °C, the digested tRNA samples were placed in a 96 well plate, mixed with 10 μL LC-MS buffer and analyzed by LC-MS/MS. Quantification was performed on an Agilent 1290 series HPLC combined with an Agilent 6470 Triple Quadrupole mass spectrometer. Nucleosides were separated using a Synergi Fusion-RP column (Synergi® 2.5 μm Fusion-RP 100 Å, 150 × 2.0 mm, Phenomenex®, Torrance, CA, USA) at a flow rate of 0.35 mL min^−1^ and a column temperature of 35 °C. Buffer A consisting of 5 mM ammoniumacetate pH 5.3 and buffer B consisting of pure acetonitrile were used as buffers. The gradient for chromatography starts with 100% buffer A for 1 min, followed by an increase to 10% buffer B over a period of 4 min. Subsequently buffer B is increased to 40% over 2 min and maintained for 1 min before switching back to 100 % buffer A over a period of 0.5 min. The column is then re-equilibrated for 2.5 min to reach starting conditions. For MS analysis the nucleosides are ionized using an ESI (electro spray ionization) source (Agilent Jetstream). The instrument was operated in positive ion mode and nucleosides were detected using a dynamic multiple reaction monitoring method.

For absolute quantification, defined concentrations of nucleosides were used as calibration. The calibration solutions ranged from 0.05 pmol to 100 pmol for canonical nucleosides and from 0.0025 pmol to 5 pmol for modified nucleosides. Ψ and D calibrations ranged from 0.01 pmol to 20 pmol. Prior to LC-MS, 1 μL of SILIS^Gen2^ (10x, (Heiss *et al*., 2021)) was co-injected with each calibration and each sample. Data analysis was performed with Agilent’s Quantitative Mass Hunter software.

### 4.6. Total Protein extraction and quantification

To obtain total protein extracts, pellets were resuspended in 100 μL of Empigen Lysis Buffer (ELB -0.5% Triton X-100, 50 mM HEPES, 250 mM NaCl, 1 mM DTT, 1 mM NaF, 2 mM EDTA, 1 mM EGTA, 1 mM PMSF, 1 mM Na_3_VO_4_ supplemented with protease inhibitors (Complete, EDTA-free, Roche, Cat. 11873580001)). Protein extracts were then sonicated for 2 cycles, at a 60% frequency, for 15 seconds each, and centrifuged 20 minutes at 200 g at 4 °C. In the end, protein in the supernatants was quantified using Pierce™ Bovine Serum Albumin (BCA) Protein Assay Kit (Thermo Fisher Scientific, Cat. 23225).

### 4.7. Insoluble Protein Extraction

To isolate the insoluble protein fractions, 100 μg of total protein was diluted in 100 μL ELB and centrifuged for 20 minutes at 16 000 g at 4 °C. The resulting pellet was solubilized in 80 μL of ELB and 20 μL of NP40 (10%). Samples were sonicated for 20 seconds and centrifuged for 20 minutes at 16000 g at 4 °C. The supernatant was removed, and 25 μL of complete ELB and 10 of μL 6x Loading Buffer were added to each pellet. After denaturation at 95 °C for 5 minutes samples were loaded into a 10% polyacrylamide gel. Gels containing total protein fractions and insoluble protein fractions were stained with blue safe (NZYTech, Cat.MB15201) for 30 minutes and revealed with the Odyssey Infrared Imaging system (Li-cor Biosciences).

### 4.8. Western Blotting

Total protein lysates were immunoblotted onto nitrocellulose membranes and incubated with the following primary antibodies: anti-ELP3 (ThermoFisher, Cat.702669, 1:200 dilution), anti-eIF2α (Cell Signalling Cat.9722, 1:1000 dilution), anti-phosphorylated eIF2α (Abcam, Cat. ab4837, 1:400 dilution), anti-puromycin antibody, clone 12D10 (Sigma-Aldrich, MABE343, 1:10000 dilution), Anti-APP (22C11, 1:1000 dilution), Anti-β Amyloid (D-11 Santacruz, Cat. Sc-374527, 1:1000) and anti-β-tubulin (Invitrogen Cat.32–2600 or Proteintech Europe, Cat.10094-1-AP, 1:1000 dilution).

### 4.9. cDNA synthesis and qPCR

cDNA was synthesized using the High-Capacity cDNA Reverse Transcription kit™ (Thermo Fisher Scientific, Cat. 4388950), according to manufacturer’s instructions. The resulting cDNA was used to perform qPCRs using the TaqMan™ Gene Expression Master Mix (Thermo Fisher Scientific, Cat.4369016) accordingly to the user guide directives. GAPDH was used as internal control to normalize gene expression levels. The reaction was carried out in the Applied Biosystems 7500 Real-Time PCR System.

### 4.10. Northern Blotting

For each sample, 5 μg of total RNA was electrophoresed on 10% polyacrylamide/urea gel. For gels containing APM, APM was added at a concentration of 20 μM in the gel mixture. RNA was transferred to a positively charged nylon membrane (Amersham Biosciences, Cat.RPN203B) in a semi-dry blotting system and auto-cross linked twice in a Stratalinker equipment (1200 mJ/cm^2^, 1 minute). Membranes were incubated in hybridization buffer (50x Denhardt’s solution, 10% SDS, 20x SSPE and miliQ water) for 2 hours, followed overnight incubation with non-radioactive probes according to their melting temperatures (5°C lower). After washing, the signal was detected using an Odyssey Infrared Imaging system (Li-cor Biosciences).

### 4.11. Statistical Analysis

Data are presented as mean with SEM and were analyzed using GraphPad Prism® software (Version 9.0).

## 5. Structured Methods

### 5.1. Reagents and Tools Table

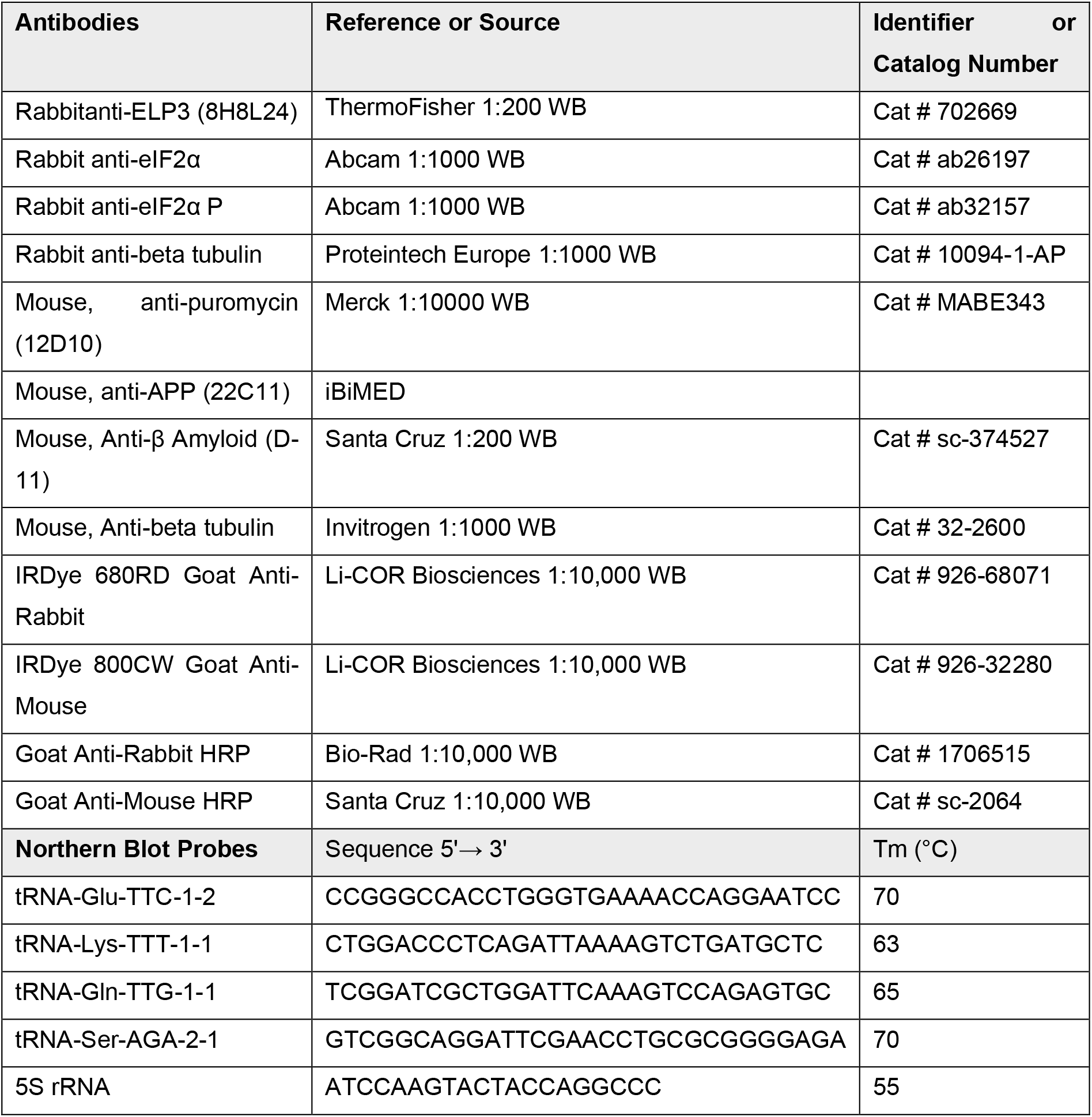

## 6. Acknowledgements

We are grateful to Professor Dora Brites (iMed.ULisbon, Faculty of Pharmacy, PT), for providing the human neuroblastoma cells: SH-SY5Y wild-type (SH-WT) and SH-SY5Y carrying the APP695 Swedish mutation (SH-SWE), to Dr Mafalda Santos (IMM, University of Lisbon), that kindly provided us the tRNA-Lys^UUU^ cloned into the pIRES2-dsRed plasmid and to Dr Hélio Albuquerque (University of Aveiro) that synthetized the APM component.

## 8. Author Contributions

Conceptualization, M.P. and A.R.S.; methodology, M.P., D.R.R., M.B., M.M., A.P.T., K.N., and A.R.S.; data curation, M.P.; writing-original draft preparation, M.P.; writing-review and editing, M.M., S.K. and A.R.S.; visualization, M.P.; supervision, A.R.S.; project administration, A.R.S.; funding acquisition, A.R.S., M.M, and S.K. All authors have read and agreed to the published version of the manuscript.

