## Supplementary material for "Amyloid pathology reduces ELP3 expression and tRNA modifications leading to impaired proteostasis in Alzheimer’s disease models": Suppl.Fig1

**A**

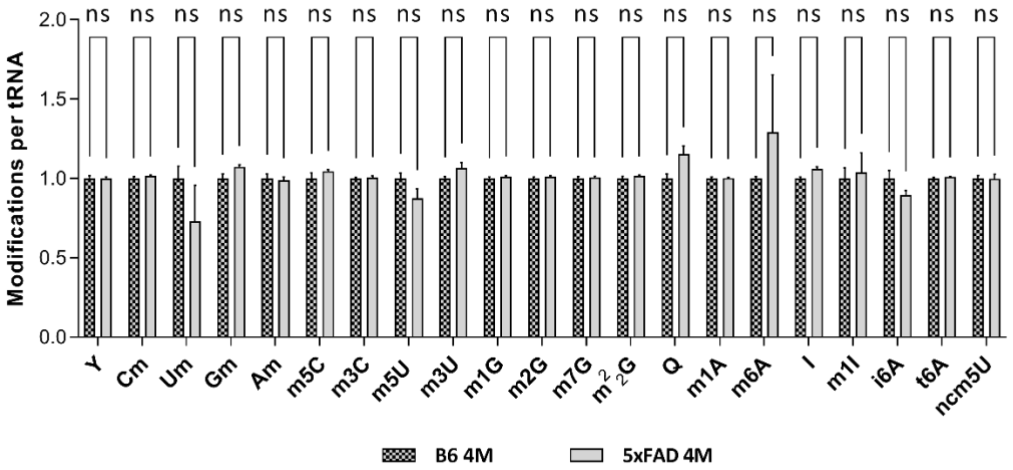


**Suppl.Fig1. Quantification of additional tRNA modifications in the hippocampus of 4 months old (4M) 5xFAD mice by LC-MS/MS.** **A)** Quantification of additional tRNA modifications in the hippocampus (Hipp) of 4M 5xFAD mice compared to B6 mice. The analysis revealed no significant alterations in the levels of tRNA modifications between the different mice models. **Data information:** data are expressed as mean SEM, n = 3 biological replicates. The non-significant (ns) p value as assessed by two-way ANOVA with the Sidak test.
