## Supplementary material for "Amyloid pathology reduces ELP3 expression and tRNA modifications leading to impaired proteostasis in Alzheimer’s disease models": Suppl.Fig2

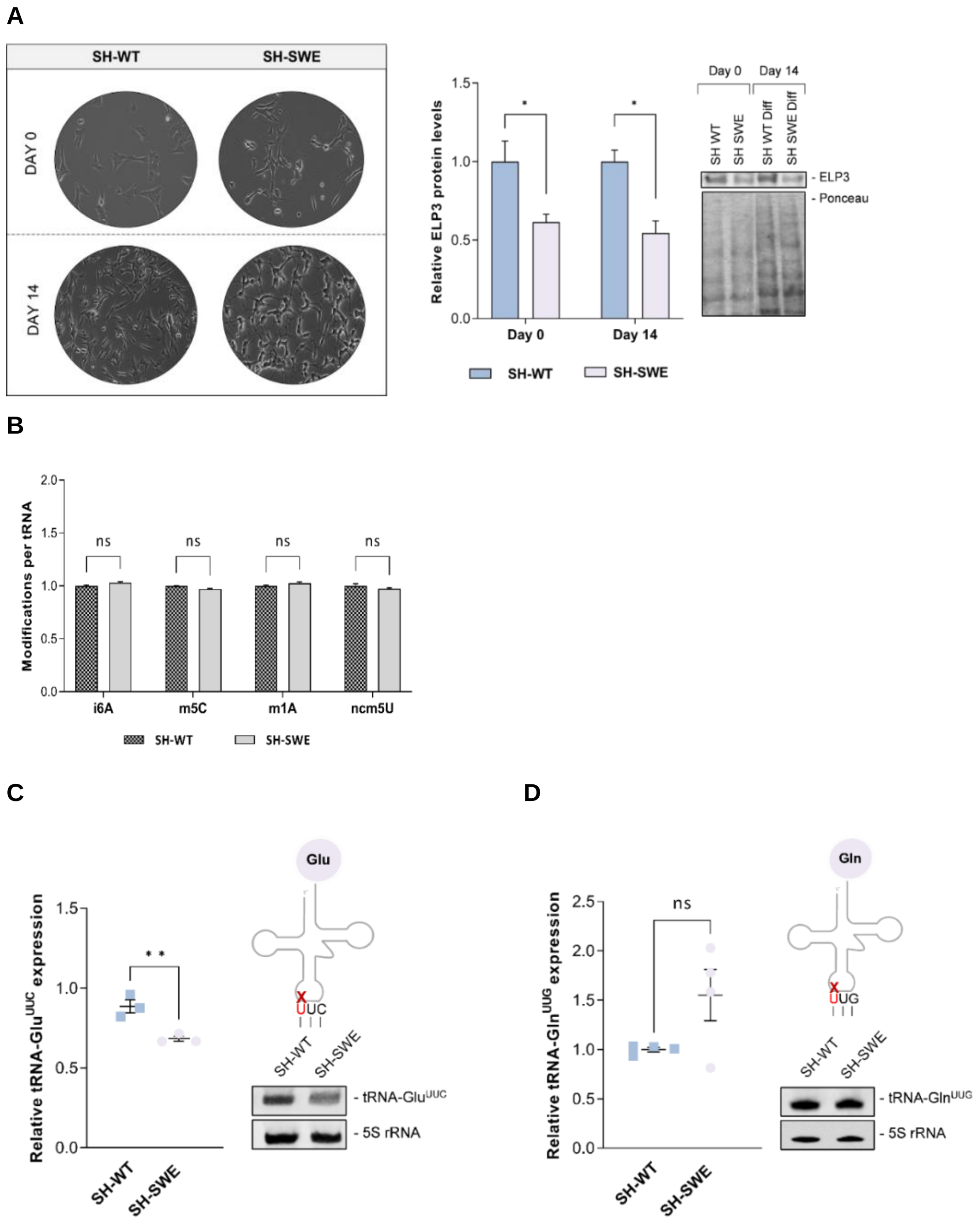


**Suppl.Fig2: ELP3 expression is decreased in SH-SWE cells. A)** Microscopy images of SH-WT and SH-SWE cells before (day 0) and after differentiation (day 14) with retinoic acid (RA: 10µM) and brain derived neurotrophic factor (BDNF: 50 ng/mL). Western blot and graphical quantification of ELP3 protein levels before (day 0) and after (day 14) SH-WT and SH-SWE cells differentiation. A significant decrease in the expression of ELP3 was observed in SH-SWE cells when compared to SH-WT cells, both at day 0 and day 14. **B)** Quantification of the i$^6$A, m$^5$C and m$^1$A and ncm$^5$U modifications per tRNA molecule in SH-SWE cells compared to SH-WT cells. The analysis revealed no significant alterations in the levels of these tRNA modifications between both cell lines. **C)** Quantification of tRNA-Glu$^{UUC}$ expression by Northern blotting. Mature tRNA-Glu$^{UUC}$ abundance is significantly decreased in SH-SWE cells, in comparison with SH-WT cells. 5S RNA was used as internal control. **D)** Quantification of tRNA-Gln$^{UUG}$ expression by Northern blot. There are no significant alterations in the abundance of the total tRNA-Gln$^{UUG}$ present in the SH-SWE cells in comparison with SH-WT cells. 5S RNA was used as internal control. **Data information**: data are expressed as mean with SEM, n = 3 biological replicates. *p-value <0.05, **p-value <0.01, and non-significant (ns) p-value as assessed by two-way ANOVA with the Sidak test (in A, B) and unpaired t test (in C, D).
