## Supplementary material for "Amyloid pathology reduces ELP3 expression and tRNA modifications leading to impaired proteostasis in Alzheimer’s disease models": Suppl.Fig3

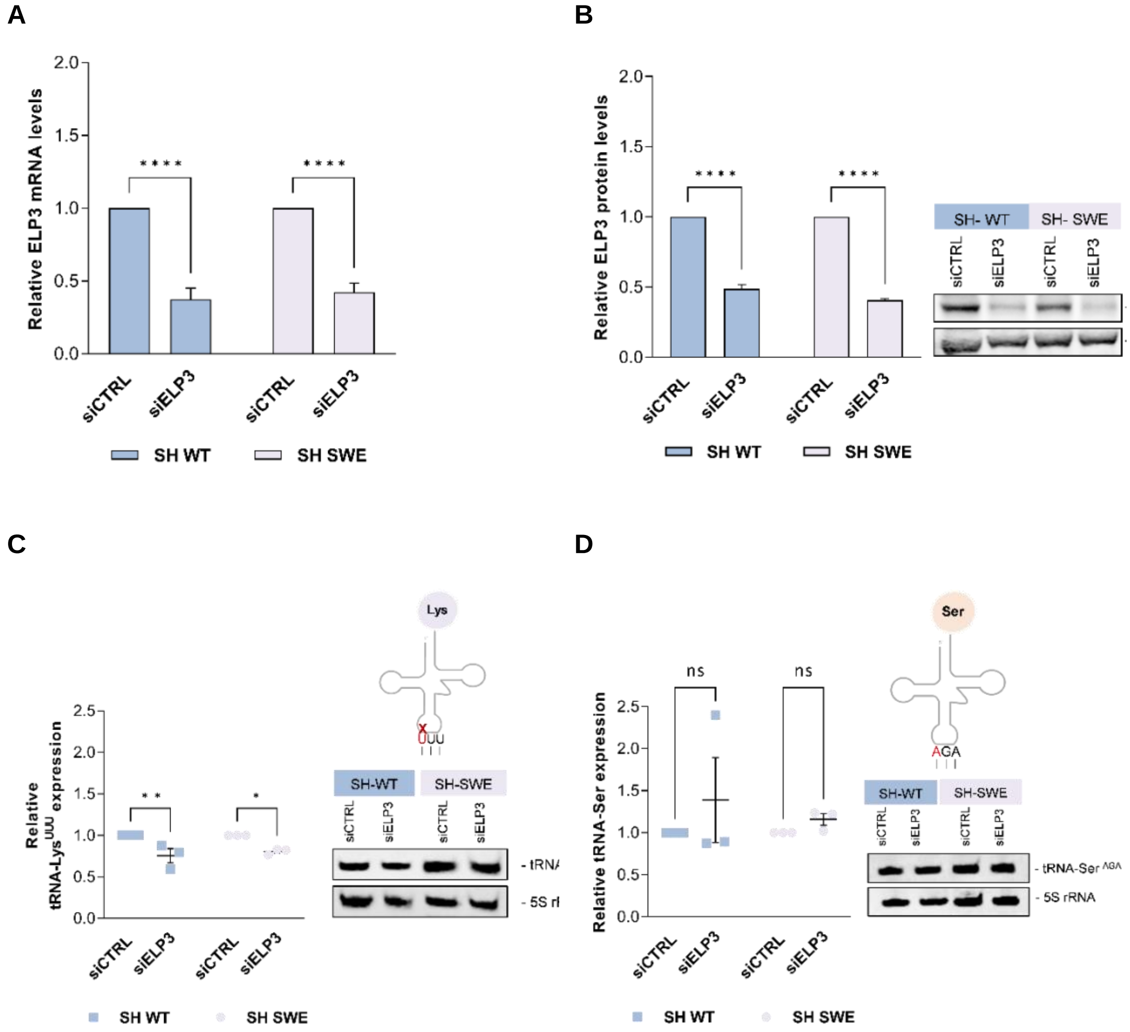


**Suppl.Fig3. Silencing efficiency of ELP3 in SH-WT and SH-SWE cells.** A) qPCR analysis of ELP3 mRNA levels in ELP3 silenced SH-WT and ELP3 silenced SH-SWE cells. A significant decrease in ELP3 mRNA levels was observed in ELP3-silenced SH-SWE (siELP3) and SH-WT (siELP3) cells when compared to cells transfected with a control siRNA (siCTRL). β-tubulin was used as an internal control. **B)** Western Blot and graphical representation of ELP3 protein levels in ELP3-silenced SH-WT and SH-SWE cells compared to siCTRL transfected cells. A significant decrease in the expression of ELP3 was observed in ELP3- silenced SH-SWE (siELP3) and ELP3-silenced SH-WT (siELP3) cells when compared to control (siCTRL) cells, in both cell lines. β-tubulin was used as an internal control. **C)** Quantification of tRNA-Lys$^{UUU}$ abundance by Northern blot after ELP3 silencing in both cell lines. A significant decrease in the abundance of total tRNA-Lys$^{UUU}$ was detected in both SH-SWE and SH-WT ELP3-silenced cells (siELP3) when compared to SH-SWE or SH-WT cells transfected with a control siRNA (siCTRL). **D)** Quantification of tRNA-Ser$^{AGA}$ abundance by Northern blot after ELP3 silencing in both cell lines. No difference in the abundance of tRNA-Ser$^{AGA}$ was detected in ELP3- silenced SH-SWE (siELP3) and ELP3-silenced SH-WT (siELP3) cells when compared to the respective control cells (siCTRL). **Data information**: data are expressed as mean with SEM, n = 3 biological replicates. *p-value <0.05, **p-value <0.01, ***p-value <0.001, ****p-value <0.0001 and non-significant (ns) p-value as assessed by two-way ANOVA with the Sidak test (A-D).
