## Supplementary material for "Amyloid pathology reduces ELP3 expression and tRNA modifications leading to impaired proteostasis in Alzheimer’s disease models": Suppl.Fig4

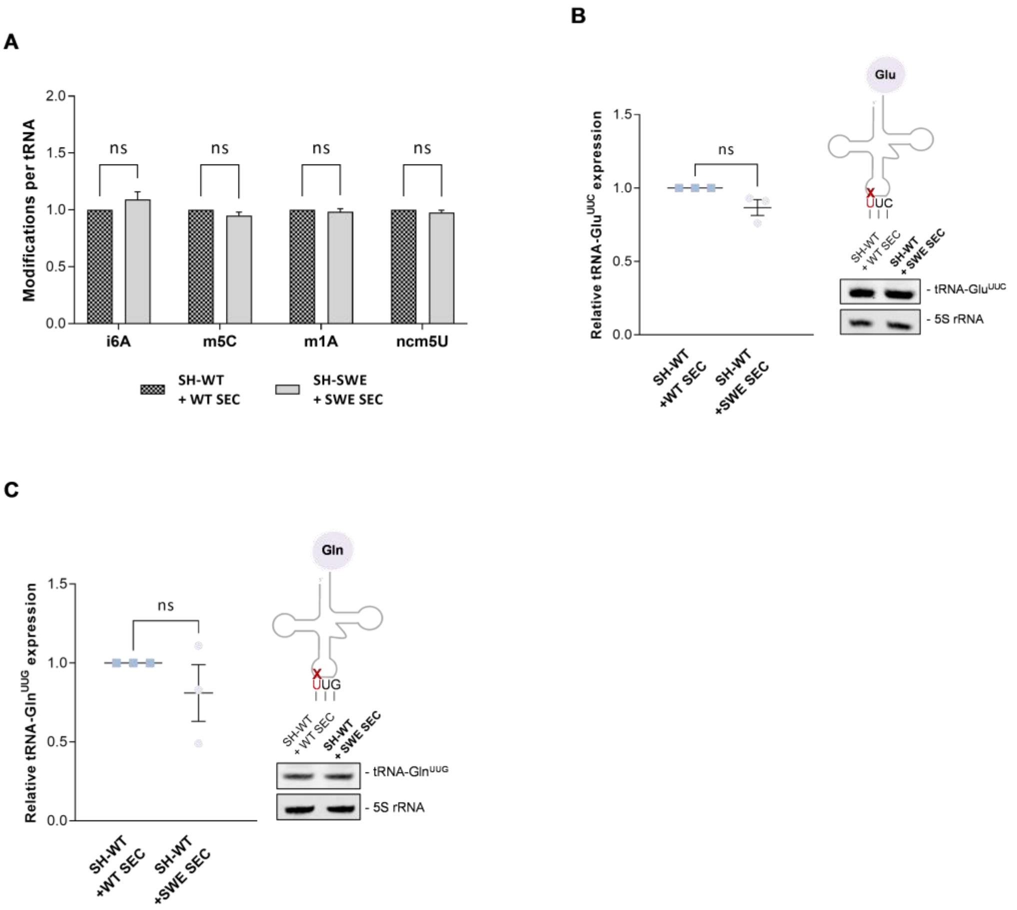


**Suppl.Fig4. Quantification of additional tRNA modifications by LC/MS-MS and tRNA-Glu$^{UUC}$ and tRNA-Gln$^{UUG}$ expression in SH-WT cells incubated with SH-SWE cells secretome. A)** Quantification of the $i^6A$, $m^5C$ and $m^1A$ and $ncm^5U$ tRNA modifications levels in SH-WT cells incubated with SH-SWE secretome (SH-WT + SWE SEC), in comparison to the control condition SH-WT cells incubated with SH-WT secretome (SH-WT + WT SEC). No significant alterations in the levels of these tRNA modifications in the SH-WT + SWE SEC cells, compared to SH-WT + WT SEC cells were detected. B) Quantification of the tRNA-Glu$^{UUC}$ levels in SH-WT cells incubated with SH-SWE secretome (SH-WT + SWE SEC), compared to the control condition SH-WT cells incubated with SH-WT secretome (SH-WT + WT SEC). No significant alterations in the abundance of total tRNA-Glu$^{UUC}$ were detected in SH-WT + SWE SEC cells, compared to SH-WT + WT SEC cells. 5S RNA was used as internal control. **C**) Quantification of tRNA-Gln$^{UUG}$ levels in SH-WT cells incubated with SH-SWE secretome (SH-WT + SWE SEC), compared to the control condition SH-WT cells incubated with SH-WT secretome (SH-WT + WT SEC). No significant alterations in the abundance of the total tRNA-Gln$^{UUG}$ present in the SH-WT + SWE SEC cells, compared to SH-WT + WT SEC cells were detected. 5S RNA was used as internal control. **Data information:** data are expressed as mean SEM, n = 3 biological replicates. The non-significant (ns) p value as assessed by two-way ANOVA with the Sidak test (in A) and unpaired t test (in B-C).
