## Supplementary material for "Amyloid pathology reduces ELP3 expression and tRNA modifications leading to impaired proteostasis in Alzheimer’s disease models": S_Table1

**Supplementary Table 1 (S_Table1)**

Levels of Log2FC and p-values of the analyzed tRNA Modifying Enzymes in different brain areas of Alzheimer´s disease patients (AD), compared to Cognitively normal older adults (CN). Abbreviations: **ACC**: Anterior Cingulate Cortex; **CBE:** Cerebellum; **DLPFC:** Dorsolateral Prefrontal Cortex; **FP**: Frontal Pole; **IFG**: Inferior Frontal Gyrus; **PCC:** Posterior Cingulate Cortex; **PHG:** Parahippocampal Gyrus; **STG**: Superior Temporal Gyrus; **TCX**: Temporal Cortex.

| | tRNAModEnz | ACC | *p-value* | CBE | *p-value* | DLPFC | *p-value* | FP | *p-value* | IFG | *p-value* | PCC | *p-value* | PHG | *p-value* | STG | *p-value* | TCX | *p-value* |
|---|---|---|---|---|---|---|---|---|---|---|---|---|---|---|---|---|---|---|---|
| Wobble Position | IKBKAP/ELP1 | 3,0E-02 | | 7,9E-02 | | -2,4E-03 | | 5,6E-03 | | -6,2E-02 | | -1,4E-03 | | -9,8E-02 | 3,0E-02 | -5,5E-02 | | -3,5E-02 | |
| | ELP2 | -4,0E-02 | | -1,0E-01 | 1,1E-03 | -8,9E-02 | 6,4E-07 | -2,9E-02 | | -5,8E-02 | 2,0E-02 | -4,9E-02 | 4,6E-02 | -2,6E-02 | | -8,0E-02 | 1,1E-03 | -1,2E-01 | 6,1E-05 |
| | **ELP3** | **-3,1E-02** | | **-3,9E-02** | | **-3,5E-02** | | **-4,1E-02** | | **-8,8E-02** | **9,4E-03** | **-5,5E-02** | **3,4E-02** | **-8,7E-02** | **4,3E-03** | **-6,0E-02** | | **-1,2E-02** | |
| | ELP4 | 2,3E-02 | | 1,2E-01 | 1,4E-02 | 1,4E-02 | | 2,7E-02 | | 1,4E-02 | | 4,3E-02 | | 3,6E-02 | | -1,2E-02 | | 2,1E-01 | 1,1E-04 |
| | ELP5 | 7,5E-02 | 5,5E-03 | 7,6E-02 | 1,6E-02 | 1,8E-02 | | -3,6E-02 | | -6,1E-03 | | 2,1E-02 | | -6,1E-02 | | 1,4E-02 | | 7,5E-02 | 1,5E-02 |
| | ELP6 | -1,9E-02 | | -3,0E-02 | | -1,2E-02 | | -7,5E-02 | 4,5E-02 | -1,3E-02 | | -5,5E-02 | | -8,9E-02 | 5,2E-03 | -5,0E-02 | | -1,0E-01 | 9,1E-03 |
| | URM1 | -1,2E-02 | | -1,2E-01 | 1,1E-02 | -5,0E-02 | 4,4E-02 | -6,7E-02 | | 5,4E-02 | | -9,6E-02 | 1,7E-03 | 3,6E-02 | | 3,1E-02 | | 6,0E-02 | |
| | CTU1 | 7,1E-02 | | 1,3E-01 | | -7,7E-03 | | -4,2E-02 | | 1,2E-02 | | 2,1E-02 | | 1,1E-01 | | 1,0E-02 | | 3,4E-01 | 5,8E-06 |
| | CTU2 | 1,8E-02 | | 1,7E-01 | 6,9E-03 | 2,3E-02 | | -1,5E-02 | | 4,3E-02 | | -9,9E-03 | | 1,5E-02 | | 4,1E-02 | | 8,3E-02 | |
| | ALKBH8 | -1,1E-01 | 3,1E-03 | 1,4E-03 | | -6,6E-02 | | -7,0E-02 | | -1,3E-01 | 3,2E-02 | -8,7E-02 | 3,8E-02 | -1,5E-01 | 5,2E-03 | -1,9E-01 | 9,8E-04 | -2,6E-01 | 7,0E-06 |
| | FTSJ1 | -7,3E-02 | 3,4E-03 | 2,6E-02 | | 8,0E-03 | | -5,2E-03 | | -7,4E-02 | 4,2E-02 | -1,3E-02 | | -7,2E-02 | 3,4E-02 | -6,7E-02 | | 1,3E-02 | |
| | QTRT1 | -6,8E-02 | | -1,9E-01 | 1,3E-02 | -1,0E-01 | | -8,2E-02 | | 3,0E-02 | | -6,6E-02 | | -6,7E-02 | | -4,2E-03 | | -1,4E-01 | |
| | NSUN2 | -9,6E-02 | 3,6E-02 | -9,6E-02 | | -1,3E-01 | 1,9E-04 | -8,0E-02 | | -5,9E-02 | | -1,1E-01 | 2,3E-02 | 2,2E-03 | | -4,7E-02 | | -3,2E-02 | |
| | TRMU | 5,8E-02 | | 1,0E-03 | | -5,5E-02 | | -5,4E-02 | | 1,7E-02 | | -1,9E-02 | | 3,8E-02 | | 2,3E-02 | | 6,3E-02 | |
| | ADAT2 | -2,9E-01 | 2,1E-07 | -3,2E-01 | 1,9E-06 | -1,9E-01 | 1,1E-05 | -1,6E-01 | 1,2E-02 | -2,2E-01 | 2,1E-04 | -2,3E-01 | 8,3E-05 | -2,8E-01 | 1,7E-07 | -2,8E-01 | 7,0E-06 | -4,3E-01 | 2,3E-08 |
| | ADAT3 | 2,0E-01 | 3,4E-02 | 4,9E-01 | 7,5E-06 | 2,0E-01 | 1,0E-02 | -2,2E-01 | | 2,2E-01 | | 2,6E-01 | 5,0E-03 | 3,9E-01 | 8,3E-04 | 9,8E-02 | | 3,2E-01 | 1,0E-02 |
| | NSUN3 | 2,1E-03 | | -5,3E-02 | | -1,3E-02 | | -2,3E-02 | | -4,1E-02 | | -4,9E-02 | | -4,3E-02 | | -9,4E-02 | | -1,5E-01 | 1,7E-03 |
| | GTPBP3 | -9,7E-02 | 3,7E-02 | -9,7E-02 | | -7,6E-02 | 2,3E-02 | -9,3E-02 | | 1,7E-02 | | -7,6E-02 | | -9,9E-03 | | -3,8E-02 | | -7,8E-02 | |
| | TRDMT1 | 3,7E-02 | | -1,9E-01 | 6,0E-03 | 3,8E-02 | | 4,7E-02 | | 8,7E-03 | | 7,0E-02 | | 8,4E-02 | 3,0E-02 | -1,9E-02 | | -7,5E-02 | |
| Outside Wobble Position | TRMT2A | 2,2E-02 | | -1,0E-01 | | -3,6E-02 | | -3,1E-02 | | 6,5E-02 | | -9,3E-03 | | 3,0E-02 | | 4,8E-02 | | 1,2E-02 | |
| | TRMT2B | 1,0E-01 | 1,5E-03 | 1,7E-01 | 6,0E-03 | 7,1E-02 | 1,4E-02 | 4,2E-02 | | 1,1E-02 | | -3,0E-02 | | 2,1E-02 | | 2,0E-02 | | 1,9E-02 | |
| | PUS1 | 1,9E-02 | | 3,5E-02 | | -8,0E-02 | 1,7E-02 | -3,7E-02 | | 8,3E-03 | | -2,5E-02 | | -7,2E-02 | | -1,9E-03 | | 5,0E-02 | |
| | PUS3 | 8,8E-02 | | 1,6E-01 | 2,2E-02 | 3,3E-03 | | -6,5E-02 | | -7,8E-02 | | 3,2E-02 | | -9,0E-02 | | -7,7E-02 | | 7,3E-02 | |
| | TRMT12 | 5,3E-02 | | 2,2E-01 | 3,4E-04 | -3,5E-02 | | -9,5E-02 | 4,0E-02 | -9,2E-02 | 3,8E-02 | -1,9E-02 | | -1,6E-01 | 3,4E-05 | -1,4E-01 | 1,4E-03 | -1,4E-01 | 2,4E-02 |
| | TRMT5 | 2,0E-03 | | -1,2E-01 | 1,4E-02 | -9,0E-03 | | 3,6E-03 | | -4,4E-02 | | 3,6E-03 | | -5,3E-02 | | -9,6E-02 | 1,1E-02 | -7,0E-02 | |
| | TRMT1 | -6,9E-02 | | 7,2E-04 | | -8,3E-02 | 3,3E-02 | -5,3E-02 | | 2,5E-02 | | -1,1E-01 | 3,8E-02 | -6,6E-02 | | 1,3E-02 | | 2,2E-02 | |
| | TRIT1 | -5,5E-02 | | -2,5E-03 | | -7,0E-02 | 8,6E-03 | -6,8E-02 | | -9,7E-02 | 1,1E-02 | -3,6E-02 | | -9,0E-02 | 7,1E-03 | -7,1E-02 | | -1,9E-02 | |
| | LCMT2 | 8,0E-02 | 1,8E-02 | -2,1E-01 | 2,5E-04 | 4,2E-03 | | -2,0E-02 | | -3,8E-02 | | 3,5E-02 | | -5,0E-02 | | -5,9E-02 | | -1,7E-02 | |
| | TRMT11 | -3,8E-03 | | -2,6E-01 | 1,1E-03 | -2,9E-02 | | -1,2E-01 | 6,6E-03 | -1,3E-01 | 1,3E-03 | -3,7E-02 | | -1,9E-01 | 3,9E-08 | -1,8E-01 | 1,4E-05 | -2,5E-01 | 5,4E-03 |
| | TRMT10A | -1,0E-01 | 2,7E-02 | -8,4E-03 | | -2,8E-02 | | -1,1E-01 | | -5,7E-03 | | -4,0E-02 | | 1,2E-01 | | -3,1E-02 | | -1,1E-01 | |
| | TRMT61A | 8,0E-02 | | -1,3E-01 | | -1,8E-02 | | -7,3E-02 | | 3,6E-02 | | 3,6E-02 | | 5,9E-02 | | 3,9E-02 | | 8,1E-02 | |
| | KTI12 | -2,7E-02 | | -2,9E-02 | | -1,4E-02 | | -2,0E-01 | | -3,6E-01 | 4,7E-04 | -9,6E-02 | | -4,4E-02 | | -1,6E-01 | | -8,1E-02 | |
| | ADAT1 | -1,6E-02 | | 9,1E-03 | | -1,3E-02 | | -5,7E-02 | | -8,0E-02 | 2,1E-02 | -5,6E-02 | | -6,0E-02 | | -8,0E-02 | 2,5E-02 | -2,0E-01 | 1,4E-04 |
| | CDK5RAP1 | -5,3E-02 | | 5,9E-02 | | -5,8E-02 | 1,6E-02 | -3,8E-02 | | -1,0E-01 | 5,3E-04 | -6,2E-02 | | -9,4E-02 | 4,0E-04 | -6,6E-02 | 3,4E-02 | -6,6E-02 | |
| | CDKAL1 | -1,1E-02 | | 1,0E-01 | 1,7E-02 | 7,7E-03 | | -1,3E-02 | | 2,4E-02 | | 9,8E-03 | | 9,8E-02 | 1,4E-03 | 3,5E-02 | | 9,5E-03 | |
| | DUS2 | -1,5E-01 | 1,1E-05 | -3,3E-01 | 4,4E-05 | -1,1E-01 | 2,7E-05 | -9,0E-02 | 3,0E-02 | -9,6E-02 | 1,3E-02 | -2,0E-01 | 4,4E-08 | -8,2E-02 | 2,2E-02 | -1,0E-01 | 1,1E-02 | -1,8E-01 | 2,1E-02 |
| | DUS1L | 7,2E-02 | | -6,0E-02 | | 1,7E-03 | | -6,5E-02 | | 4,8E-02 | | 3,8E-02 | | 2,9E-02 | | 3,5E-02 | | 3,3E-02 | |
| | DUS3L | 2,6E-02 | | -1,0E-01 | | -9,9E-03 | | -1,3E-01 | 3,7E-02 | 1,1E-01 | 3,9E-02 | 1,9E-02 | | 3,7E-02 | | -2,9E-02 | | -1,2E-01 | |
| | DUS4L | -1,6E-01 | 1,2E-04 | -4,4E-01 | 4,0E-07 | -1,1E-01 | 2,8E-03 | -7,8E-02 | | -1,4E-01 | 8,9E-03 | -9,7E-02 | 4,1E-02 | -1,5E-01 | 1,5E-03 | -1,4E-01 | 1,2E-02 | -3,3E-01 | 2,4E-05 |
| | METTL1 | -2,0E-01 | 3,7E-03 | -1,5E-01 | 9,1E-02 | -2,0E-01 | 3,7E-04 | -6,8E-02 | | -2,2E-01 | 2,3E-02 | -2,4E-01 | 3,2E-03 | -7,9E-02 | | -8,7E-02 | | -7,7E-02 | |
| | METTL2A | -2,6E-02 | | -2,0E-02 | | -4,6E-02 | | -5,4E-02 | | -6,8E-02 | | -9,7E-02 | 9,9E-04 | -7,4E-02 | 2,5E-02 | -5,5E-02 | | -7,5E-02 | 2,1E-02 |
| | METTL2B | -1,8E-02 | | -6,3E-02 | | -3,4E-02 | | -2,1E-02 | | -4,5E-02 | | -8,2E-02 | | -3,8E-03 | | -1,2E-02 | | -6,6E-02 | |
| | METTL6 | -8,6E-02 | 3,4E-04 | -1,8E-02 | | -4,5E-02 | 3,2E-02 | -7,7E-02 | 1,7E-02 | -9,6E-02 | 1,5E-03 | -8,1E-02 | 1,6E-03 | -9,1E-02 | 7,1E-04 | -9,6E-02 | 1,7E-03 | -2,0E-01 | 6,3E-10 |
| | NSUN6 | -2,3E-02 | | -9,5E-03 | | -8,5E-02 | 9,0E-04 | -7,7E-02 | | -1,3E-01 | 3,9E-04 | -5,7E-02 | | -6,0E-02 | | -5,9E-02 | | -1,2E-01 | 9,7E-03 |
| | OSGEP | -5,2E-02 | | -3,3E-01 | 2,3E-04 | -1,3E-01 | 2,7E-03 | -1,8E-01 | 4,0E-03 | -1,2E-01 | 4,0E-02 | -1,0E-01 | | -9,6E-02 | | -1,3E-01 | 3,6E-02 | -2,7E-01 | 2,9E-03 |
| | PUS10 | -9,5E-02 | 2,4E-02 | -6,0E-02 | | -1,2E-01 | 4,4E-04 | -1,1E-02 | | -1,8E-02 | | -9,0E-02 | 3,4E-02 | -8,1E-02 | | -6,5E-02 | | -2,1E-01 | 5,3E-04 |
| | PUS7 | -3,7E-03 | | 9,6E-02 | | -1,7E-02 | | -4,0E-02 | | -1,3E-01 | 9,1E-04 | -6,0E-02 | | -8,8E-02 | 1,4E-02 | -7,1E-02 | | 5,9E-03 | |
| | THG1L | 8,6E-02 | | -4,7E-03 | | 5,3E-03 | | -1,1E-01 | 3,5E-02 | -8,8E-02 | | -5,6E-03 | | -2,9E-02 | | -2,8E-02 | | -9,8E-02 | 3,8E-02 |
| | TRMO | 2,6E-02 | | -1,2E-03 | | 3,8E-03 | | -8,2E-02 | | -2,1E-02 | | 3,6E-02 | | 4,8E-02 | | 6,6E-03 | | -3,0E-02 | |
| | TRMT10B | 1,8E-02 | | 1,6E-01 | 6,8E-05 | -1,6E-02 | | 6,9E-02 | | 8,6E-02 | 1,8E-02 | 1,4E-02 | | 1,7E-01 | 1,4E-07 | 1,2E-01 | 9,3E-04 | 3,1E-02 | |
| | TRMT10C | 2,6E-02 | | -2,1E-02 | | 6,9E-03 | | -1,7E-01 | 1,7E-02 | -2,4E-01 | 3,8E-04 | -7,9E-02 | | -3,5E-01 | 8,5E-09 | -2,6E-01 | 1,8E-04 | -1,2E-01 | 3,9E-02 |
| | TRMT13 | 1,7E-03 | | -9,5E-03 | | 1,4E-02 | | 2,5E-02 | | 2,2E-02 | | 1,4E-02 | | 1,1E-01 | 1,5E-02 | -2,5E-02 | | -1,2E-01 | |
| | TRMT44 | 8,7E-02 | 1,0E-02 | 1,5E-01 | 2,8E-04 | -5,4E-02 | | 3,0E-02 | | 3,4E-02 | | 6,3E-02 | | -6,4E-03 | | 7,9E-02 | | -8,4E-02 | |
| | TRMT61B | -2,7E-02 | | 1,7E-01 | 4,9E-03 | -3,3E-02 | | -7,3E-02 | | -7,2E-02 | | -5,4E-02 | | -1,1E-01 | 1,8E-03 | -5,8E-02 | | -1,6E-01 | 1,4E-02 |
| | TYW3 | 9,5E-02 | 1,7E-02 | 8,8E-02 | | 1,0E-02 | | 1,1E-02 | | -5,1E-02 | | 5,5E-02 | | -5,5E-02 | | -6,7E-02 | | 2,5E-02 | |
| | TRMT6 | -5,0E-02 | | 2,6E-01 | 2,0E-05 | 1,7E-02 | | -4,2E-02 | | -1,5E-01 | 1,1E-02 | -7,3E-02 | | -1,6E-01 | 3,5E-03 | -1,4E-01 | 2,4E-02 | -8,9E-02 | |
