## Supplementary material for "Amyloid pathology reduces ELP3 expression and tRNA modifications leading to impaired proteostasis in Alzheimer’s disease models": S_Table2

**Supplementary Table 2 (S_Table2)**

Convergent functional genomic (CFG) ranking for target genes associated. Information collected from www.alzdata.org. Abbreviations: expression quantitative trait loci (eQTL); genome-wide association studies (GWAS); protein-protein interaction (PPI), and differentially expressed genes (DEG).

| | Gene | eQTL | GWAS | PPI | Early_DEG | Pathology cor (abeta) | Pathology cor (tau) | CFG |
|---|---|---|---|---|---|---|---|---|
| Wobble Position | IKBKAP/ELP1 | 1 | 0 | - | yes | -0.391,** | -0.088,ns | 3 |
| | ELP2 | 2 | 0 | - | NA | NA | NA | 1 |
| | ELP3 | 1 | 1 | - | yes | -0.456,** | -0.602,* | 4 |
| | ELP4 | 3 | 0 | - | NA | NA | NA | 1 |
| | ELP5 | NA | 0 | - | NA | NA | NA | 0 |
| | ELP6 | NA | 0 | - | NA | NA | NA | 0 |
| | URM1 | 2 | NA | - | NA | NA | NA | 1 |
| | CTU1 | 0 | 0 | - | NA | NA | NA | 0 |
| | CTU2 | 0 | 0 | - | NA | NA | NA | 0 |
| | ALKBH8 | 0 | 3 | - | NA | NA | NA | 1 |
| | FTSJ1 | 1 | NA | - | NA | NA | NA | 1 |
| | QTRT1 | 0 | 0 | - | yes | -0.173,ns | 0.261,ns | 1 |
| | NSUN2 | 0 | 0 | - | NA | -0.093,ns | -0.427,ns | 0 |
| | TRMU | 1 | 0 | - | NA | NA | NA | 1 |
| | ADAT2 | 0 | 0 | - | NA | NA | NA | 0 |
| | ADAT3 | 6 | 0 | - | NA | NA | NA | 1 |
| | NSUN3 | 1 | 0 | - | NA | -0.055,ns | -0.102,ns | 1 |
| | GTPBP3 | 0 | 0 | - | NA | 0.079,ns | -0.405,ns | 0 |
| | TRDMT1 | 3 | 0 | - | NA | NA | NA | 1 |
| Outside Wobble Position | TRMT2A | 1 | 0 | - | NA | NA | NA | 1 |
| | TRMT2B | 0 | NA | - | NA | NA | NA | 0 |
| | PUS1 | 1 | 0 | - | NA | -0.068,ns | -0.012,ns | 1 |
| | PUS3 | NA | 0 | - | NA | -0.209,ns | -0.195,ns | 0 |
| | TRMT12 | 3 | 0 | - | yes | -0.222,ns | -0.721,** | 3 |
| | TRMT5 | 0 | 0 | - | NA | NA | NA | 0 |
| | TRMT1 | 0 | 0 | - | NA | 0.160,ns | -0.052,ns | 0 |
| | TRIT1 | 1 | 29 | NA | NA | 0.062,ns | 0.283,ns | 2 |
| | LCMT2 | 6 | 0 | - | NA | NA | NA | 1 |
| | TRMT11 | 0 | 0 | - | NA | NA | NA | 0 |
| | TRMT10A | NA | 0 | - | NA | NA | NA | 0 |
| | TRMT61A | 3 | 1 | - | NA | NA | NA | 2 |
| | KTI12 | NA | 0 | - | NA | NA | NA | 0 |
| | ADAT1 | 2 | 0 | - | yes | -0.106,ns | -0.545,* | 3 |
| | CDK5RAP1 | 1 | 0 | - | yes | -0.188,ns | 0.120,ns | 2 |
| | CDKAL1 | 1 | 3 | - | NA | -0.349,* | -0.046,ns | 3 |
| | DUS2 | NA | 0 | - | NA | NA | NA | 0 |
| | DUS1L | 1 | 0 | - | NA | NA | NA | 1 |
| | DUS3L | 3 | 0 | - | NA | -0.066,ns | -0.708,** | 2 |
| | DUS4L | 0 | 0 | - | NA | NA | NA | 0 |
| | METTL1 | 0 | 0 | - | NA | -0.465,** | -0.327,ns | 1 |
| | METTL2A | 1 | 0 | PSEN1,PSEN2 | NA | NA | NA | 2 |
| | METTL2B | 0 | 0 | PSEN1,PSEN2 | NA | NA | NA | 1 |
| | METTL6 | 2 | 0 | - | NA | NA | NA | 1 |
| | NSUN6 | 0 | 0 | - | NA | NA | NA | 0 |
| | OSGEP | 0 | 0 | - | NA | 0.221,ns | 0.619,* | 1 |
| | PUS10 | 1 | 0 | - | NA | NA | NA | 1 |
| | PUS7 | 1 | 0 | - | NA | NA | NA | 1 |
| | THG1L | 2 | 0 | - | NA | -0.348,* | -0.079,ns | 2 |
| | TRMO | NA | NA | NA | NA | NA | NA | 0 |
| | TRMT10B | NA | 0 | - | NA | NA | NA | 0 |
| | TRMT10C | NA | 0 | - | NA | NA | NA | 0 |
| | TRMT13 | NA | 0 | - | NA | NA | NA | 0 |
| | TRMT44 | NA | 0 | - | NA | NA | NA | 0 |
| | TRMT61B | 1 | 0 | - | NA | NA | NA | 1 |
| | TYW3 | 0 | 0 | - | NA | NA | NA | 0 |
| | TRMT6 | 0 | 0 | - | NA | NA | NA | 0 |
